# Defining the molecular interaction between influenza hemagglutinin and MHC-II

**DOI:** 10.64898/2026.07.17.738765

**Authors:** Bernadeta Dadonaite, Annie Dosey, Jenny J. Ahn, Timothy C. Yu, Sara A. Sunshine, Ariana G. Farrell, Neil P. King, Jesse D. Bloom

## Abstract

The hemagglutinin (HA) of some influenza viruses can bind MHC-II, but how these proteins interact is unclear. Here we demonstrate that diverse H5 HAs can use avian or human MHC-II to enter cells. We define the molecular interface by using pseudovirus deep mutational scanning to measure how mutations to both H5 HA and an avian MHC-II affect the interaction between the two proteins. We leverage identification of mutations to HA and MHC-II that increase their binding to each other to determine a 3.4 Å cryo-EM structure of the complex. The structure shows that HA binds to MHC-II using a surface on the globular head distinct from the sialic acid receptor-binding pocket. The recognition surface on MHC-II consists of a portion of the alpha chain near the peptide-binding groove. We further show that H7 HA interacts with MHC-II similarly to H5 HA, and that H1, H2, and H9 HAs also bind avian or human MHC-II.

**Significance:** Influenza viruses are a public health burden because they cause seasonal epidemics and have led to multiple pandemics in the past. Influenza hemagglutinin has long been known to use sialic acid as a receptor for entering cells, but recently some hemagglutinins have been shown to also use MHC-II proteins for entry. Here we combine pseudovirus deep mutational scanning and cryo-EM to define the functional and structural basis of the interaction between an H5 hemagglutinin and MHC-II. We also demonstrate that hemagglutinins from multiple other influenza subtypes can utilize MHC-II to enter cells. Overall, our work uses a novel combination of experimental approaches to define the molecular basis of a previously poorly understood virus-host interaction.

## Introduction

Influenza hemagglutinin (HA) mediates viral entry by binding to host-cell receptors. For most influenza viruses, receptor binding is mediated by interactions between HA and sialic acids^1,2^, but recent work has shown that some HAs also or instead can use major histocompatibility complex class II (MHC-II) proteins as entry receptors^3–7^. Bat influenza virus H17 and H18 HAs and the avian influenza virus H19 HA do not bind sialic acids but can use MHC-II for cell entry^3,4,7^, whereas some H2 and H3 HAs can use both sialic acids and MHC-II for cell entry^5,6^. However, the molecular basis and biological relevance of HA’s interaction with MHC-II remains poorly understood. No structure of HA bound to MHC-II is available, the HA surface that binds MHC-II has not been fully defined, and the MHC-II residues that interact with HA remain unknown although there is evidence that the MHC-II alpha chain is a key determinant^3,8^.

Here we define the molecular binding interface between HA and an avian MHC-II by using pseudovirus deep mutational scanning^9,10^ to measure how nearly all amino-acid mutations to both H5 HA and MHC-II affect the interaction between the two proteins. We leverage identification of mutations that increase HA binding to a tufted duck MHC-II to determine a cryogenic electron microscopy (cryo-EM) structure of HA complexed with MHC-II. We further show that MHC-II binding is detectable across HAs from many strains from multiple influenza subtypes, including H1, H2, H5, H7, and H9.

## Results

### Diverse H5 HAs can use MHC-II for cell entry

To experimentally separate the roles of sialic acid and MHC-II in mediating cell entry, we used 293 cells that had been engineered to eliminate most sialic acid expression by knocking-out all ST3Gal and ST6Gal sialyltransferases^11,12^, hereafter referred to as “noSA” cells (**Fig. 1a**). Note 293 cells do not normally express any MHC-II^13^ (**Supplementary Fig. 1a**). We engineered these noSA cells to express avian or human MHC-II (**Fig. 1a** and **Supplementary Fig. 1a-b**). MHC-II alleles are highly variable between and within species^14,15^ (**Supplementary Fig. 1b**). For avian MHC-II, we used a DR-like isotype from a tufted duck (*Aythya fuligula*), which is a natural host for H5 influenza viruses^16,17^. For human MHC-II, we used a 15:03 DRB1 allele of the beta chain paired with a 01:01 DRA1 allele of the alpha chain. Cell-surface MHC-II expression in the 293-noSA cells engineered to express human MHC-II was comparable to its endogenous expression in human B-cell lines or an interferon-gamma treated human lung epithelial cell line (**Supplementary Fig. 1a**). We are unable to similarly quantify MHC-II expression in the 293-noSA cells engineered to express tufted duck MHC-II due to a lack of antibodies for tufted duck MHC-II and comparator cell lines from this species. As a control we also used 293 cells engineered to primarily express α2,3-linked sialic acids (referred to as “SA23” cells) or α2,6-linked sialic acids (“SA26” cells).

**Figure 1.**
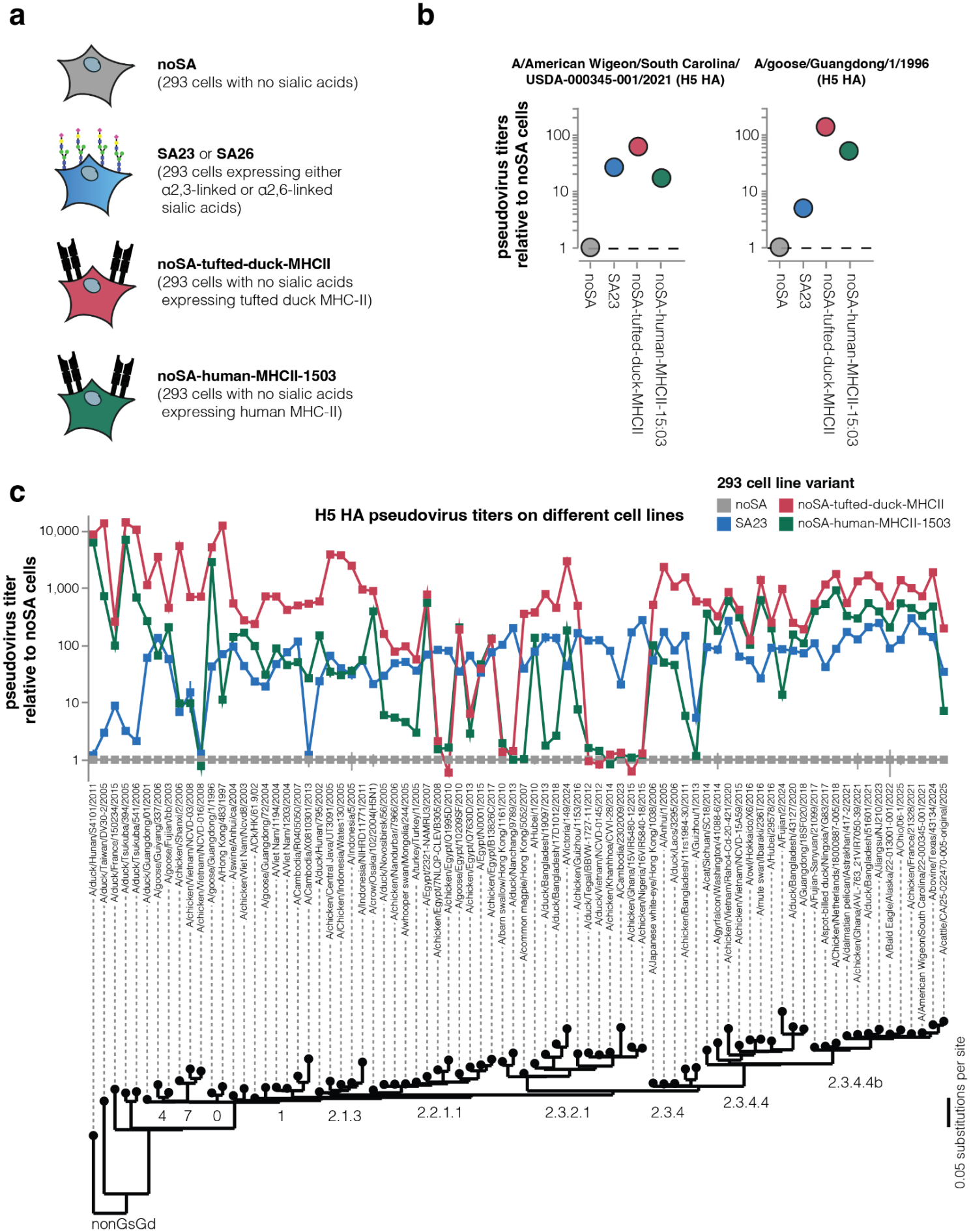
Many H5 influenza HAs can use MHC-II for cell entry. **a**. We engineered 293 cells lacking sialic acid (noSA cells) to express tufted duck or human MHC-II, and as controls used 293 cells expressing α2-3-linked or α2-6-linked sialic acids. **b**. Titers of lentiviral particles pseudotyped with the indicated H5 HA on cells expressing α2-3-linked sialic acids (SA23) or different MHC-II alleles relative to titers on no-sialic acid (noSA) cells. **c**. Titers on the different cell lines for pseudoviruses with HAs from 80 different H5 influenza strains. The phylogenetic clade is indicated by the labels on the tree aligned to the x-axis. See https://dms-vep.org/Flu-H5N1-American-Wigeon-2021-HA-tufted-duck-MHCII-DMS/entry_titers.html for an interactive version of this plot. For the same plot showing raw titers rather than titers normalized to the noSA cells, see **Supplementary Fig. 2**.

We quantified infection of the cells by lentiviral particles pseudotyped with HA from two H5N1 strains: the clade 2.3.4.4b A/American Wigeon/South Carolina/USDA-000345-001/2021 strain or the A/goose/Guangdong/1/1996 strain that is representative of early highly pathogenic H5 avian influenza viruses^18^ (**Supplementary Fig. 1c**). As expected, pseudoviruses with HAs from both strains infected the noSA cells poorly (**Fig. 1b** and **Supplementary Fig. 1d**). Expression of tufted duck MHC-II in the noSA cells greatly enhanced infection by pseudoviruses with either H5 HA (**Fig. 1b**). Pseudoviruses with either HA could also infect noSA cells expressing human MHC-II, although entry in the cells expressing human MHC-II was less efficient than in cells expressing tufted duck MHC-II (**Fig. 1b**). Expressing tufted duck MHC-II in 293T cells, which also express both α2,3- and α2,6-linked sialic acids^19,20^, only modestly increased pseudovirus titers relative to unmodified 293T cells—and only minimally increased pseudovirus titers relative to 293T cells when the neuraminidase protein (which cleaves sialic acids) was inhibited by oseltamivir (**Supplementary Fig. 1e**). The fact that expression of tufted duck MHC-II greatly increases titers in cells that lack sialic acid but only modestly increases titers in sialic acid expressing cells suggests that the two receptors mediate infection in a largely redundant fashion.

To assess how widespread MHC-II mediated entry is among H5 HAs, we chose HAs from 80 strains spanning the H5 HA phylogenetic tree, pseudotyped them on lentiviral particles, and measured titers on the different cell lines (**Fig. 1c, Supplementary Fig. 2**). In these pseudoviruses we paired the same NA (A/turkey/Indiana/22-003707-003/2022) with all HAs, as pseudovirus entry titers do not depend on which NA was used (**Supplementary Fig. 1f**), consistent with NA being involved in viral egress rather than entry^21^. MHC-II expression markedly increased pseudovirus entry for most H5 HAs. Most HAs could also use human MHC-II for cell entry, although usually with substantially lower efficiency than the tufted duck MHC-II. A minority of H5 HAs in the 2.2.1.1 and 2.3.2.1 clades have lost the ability to use tufted duck MHC-II for cell entry. Interestingly, a few HAs, like those from the low pathogenicity non-GsGd lineage, had very poor pseudovirus titers in sialic acid expressing cells but high titers in both avian and human MHC-II expressing cells.

To confirm that MHC-II mediated entry occurs in the context of influenza virions and not only in a pseudovirus context, we generated conditionally replicative influenza virions with HA and NA from A/American Wigeon/South Carolina/USDA-000345-001/2021 strain (**Supplementary Fig. 3a**). These virions do not express the essential PB1 viral polymerase protein, meaning that they can only replicate in cells constitutively expressing PB1, making them safe to use at biosafety-level-2^22^. The single-round entry titers of these influenza virions were ∼25-fold higher on noSA cells expressing tufted duck MHC-II compared to noSA cells lacking MHC-II, demonstrating that MHC-II can enable H5 HA mediated cell entry for influenza virions as well as pseudoviruses (**Supplementary Fig. 3b**).

### Effects of HA mutations on tufted duck MHC-II mediated cell entry

To define the HA sequence determinants of MHC-II mediated cell entry, we used pseudovirus deep mutational scanning^9^ to measure how mutations to the HA of the A/American Wigeon/South Carolina/USDA-000345-001/2021 (H5N1) strain affect entry into 293 cells expressing either tufted duck MHC-II or sialic acids. These experiments used previously described^10^ HA pseudovirus libraries that contain nearly all single amino-acid mutations; note that these pseudoviruses encode in their genomes no viral proteins other than HA and so can only undergo a single round of cell entry, providing a way to study HA mutations at biosafety level 2 outside the context of replicative influenza virus.

Although most HA mutations had similar effects on entry in sialic acid or tufted duck MHC-II expressing cells, at a subset of sites mutations had very different effects on entry between the cell lines (**Fig. 2a, Supplementary Fig. 4** and **5**). The mutations with different effects clustered in two regions of HA. In the sialic acid receptor-binding pocket, mutations were often more deleterious for entry in sialic acid expressing cells than MHC-II expressing cells. In contrast, mutations tended to be more deleterious for entry in MHC-II expressing versus sialic acid expressing cells in a patch of sites on the HA globular head below the receptor binding pocket (**Fig. 2a,b, Supplementary Fig. 4** and **5**, and interactive plots at https://dms-vep.org/Flu-H-5-N1American-Wigeon-2021-HA-tufted-duck-MHCII-DMS/entry.html). Specifically, mutations at sites 77-80, 120, and 141-150 (H3 numbering) impaired entry in tufted duck MHC-II expressing cells more than in sialic acids expressing cells. These sites define a potential MHC-II binding interface on the HA head domain (**Fig. 2b**) and suggest that largely distinct regions of HA bind sialic acids versus MHC-II.

**Figure 2.**
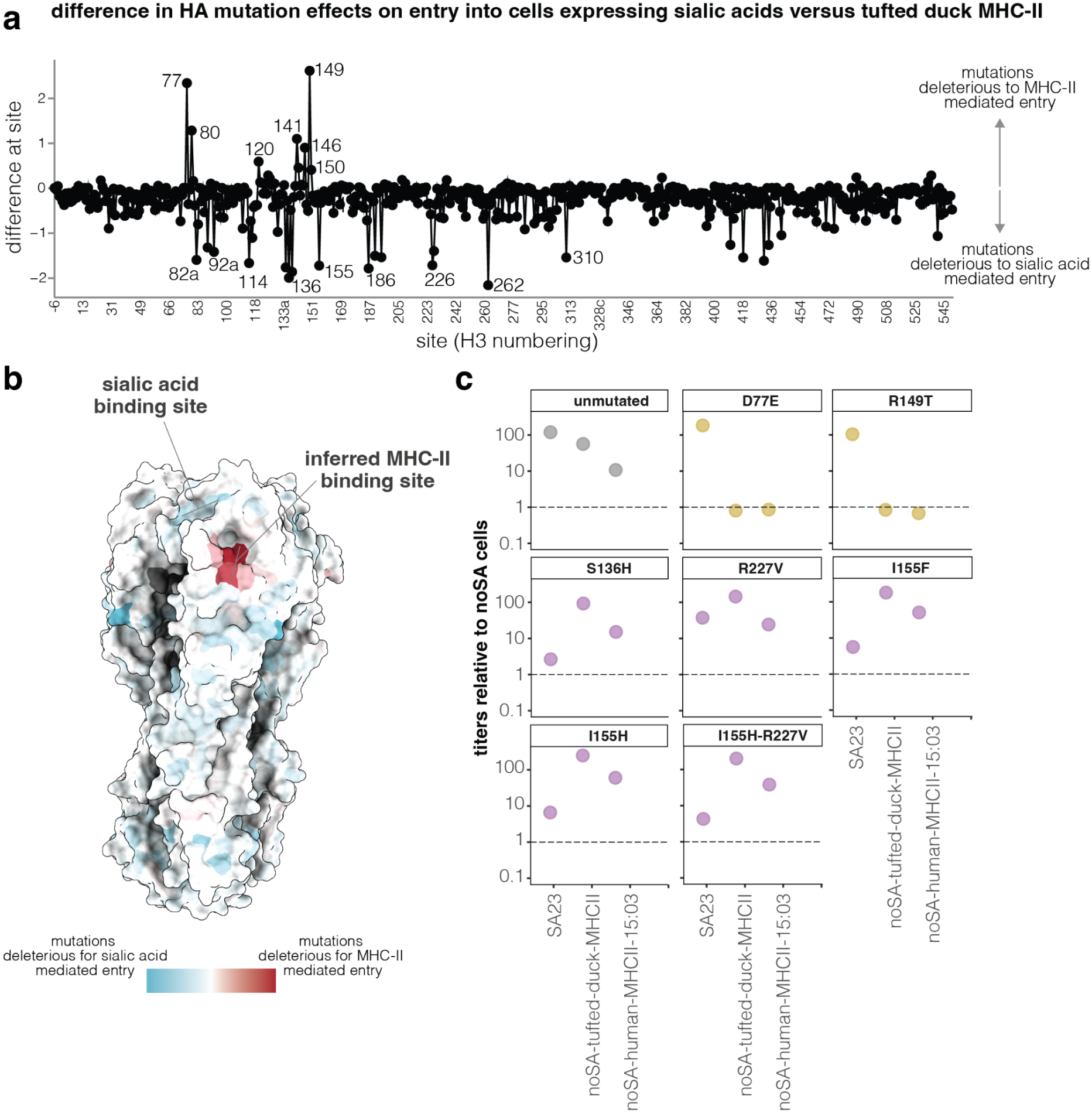
Effects of H5 HA mutations on MHC-II-mediated cell entry. **a**. Summed difference of effects of all mutations at each HA site on pseudovirus entry in a mixture of SA23 and SA26 cells versus noSA-tufted-duck-MHCII cells. Positive values indicate mutations are more deleterious for entry in noSA-tufted-duck-MHCII cells, negative values indicate mutations are more deleterious on entry in sialic acid expressing cells. See https://dms-vep.org/Flu-H5N1-American-Wigeon-2021-HA-tufted-duck-MHCII-DMS/entry.html for an interactive version of this plot that also shows effects of individual mutations. **b**. Structure of H5 HA (PDB: 4KWM) colored by the effects of mutations on sialic acid versus tufted duck MHC-II mediated cell entry; red indicates sites where mutations are more deleterious for MHC-II mediated entry, and blue indicates sites where mutations are more deleterious for sialic acid mediated entry. **c**. Validation assays showing that key HA mutations identified in the deep mutational scanning differentially affect pseudovirus entry in cells expressing sialic acids versus MHC-II. Titers are reported as fold increase relative to noSA cells. Sites are labeled using H3 numbering here and throughout this paper.

To validate the deep mutational scanning, we generated pseudoviruses with HA loss-of-function mutations for only tufted duck MHC-II or only sialic acid mediated cell entry (**Fig. 2c**). As expected from the deep mutational scanning, D77E and R149T reduced entry in tufted duck and human MHC-II expressing cells to levels comparable to that in the noSA cells while minimally impacting on entry in sialic acid expressing cells. In contrast, mutations S136H, R227V, I155F, and I155H greatly reduced pseudovirus entry in sialic acid expressing cells but did not impact entry in MHC-II expressing cells.

To confirm that these HA mutations also caused a loss-of-function for MHC-II or sialic acid mediated entry in the context of influenza virions, we introduced them into conditionally replicative influenza virions^22^ with HA and NA from A/American Wigeon/South Carolina/USDA-000345-001/2021 (H5N1) (**Supplementary Fig. 3a**). As expected, viruses with the MHC-II loss-of-function mutations (D77E or R149T) could be efficiently generated and grown in sialic acid expressing cells, and had high single-cycle entry titers in cells expressing both sialic acid and MHC-II but low entry titers in cells expressing only MHC-II and no sialic acid (Supplementary Fig. 6a,b). Also as expected, viruses with sialic acid loss-of-function mutations (S136H and I155H) could not be efficiently generated in sialic acid expressing cells (**Supplementary Fig. 6a**). However, efforts to produce viruses with the sialic acid loss-of-function mutations in MHC-II expressing cells were confounded by the fact that any virus with a HA that retained the ability to bind MHC-II formed foci when grown in MHC-II expressing cells but did not spread efficiently across the cells (**Supplementary Fig. 6a**). This pattern is reminiscent of what is observed when neuraminidase-deficient influenza viruses are grown in sialic acid expressing cells^23,24^, since when progeny virions bind to the producing cell they can only infect spatially adjacent neighbor cells (normally neuraminidase cleaves the sialic acid that HA binds on the producing cell). Our data suggest that HA can bind cell-surface MHC-II on the producing cell, hindering virion release and limiting infection to foci of spatially adjacent cells (**Supplementary Fig. 6a**). Consistent with this hypothesis, the only viruses that spread efficiently in dual sialic acid and MHC-II expressing cells were those with the MHC-II loss-of-function mutations (**Supplementary Fig. 6a,b**). It is unclear whether similar trapping of progeny virions by binding to cell-surface MHC-II occurs in authentic infections: MHC-II expression in the cell lines used in our experiments may be higher than endogenous expression, and other viral proteins expressed by authentic H5N1 could conceivably modulate MHC-II expression of infected cells^25^.

### Effects of HA mutations on binding to tufted duck MHC-II

To directly measure how HA mutations affect MHC-II binding, we leveraged the fact that pseudovirus neutralization by soluble protein receptor is proportional to the strength of binding between that receptor and the corresponding viral entry protein^26,27^. To increase the potency of the MHC-II neutralization by putting it in a multivalent format that increases binding avidity, we generated MHC-II coated virus-like particles (MHCII-VLPs) by cotransfecting cells with plasmids encoding lentiviral Gag and tufted duck MHC-II (**Supplementary Fig. 7a**). These MHCII-VLPs potently neutralized H5 HA pseudoviruses, whereas bald VLPs (produced by transfecting cells with only Gag-expressing plasmids) showed much weaker neutralization, indicating that HA binding to the MHC-II on the VLPs was the major driver of the pseudovirus neutralization (**Supplementary Fig. 7b**). Note however that because Gag forms VLPs enveloped by patches of the cell plasma membrane^28^, these VLPs likely also contain other cell-surface sialylated glycoproteins that may bind to pseudovirus HA via its sialic acid binding pocket.

We used the H5 HA deep mutational scanning libraries to measure how HA mutations that maintain at least modest cell entry function affect binding to tufted duck MHCII-VLPs as assessed by pseudovirus neutralization (**Fig. 3a-c, Supplementary Fig. 7c**, and interactive plots at https://dms-vep.org/Flu-H5N1-American-Wigeon-2021-HA-tufted-duck-MHCII-DMS/binding.html). The mutations that affect MHCII-VLP binding are all in the globular head of HA (**Fig. 3a-c**), with most mutations that strongly reduce MHCII-VLP binding at sites in the same region below sialic acid binding pocket where mutations specifically impair entry in MHC-II versus sialic acid expressing cells (compare **Fig. 3c** and **Fig. 2b**). Specifically, the mutations that most reduced MHCII-VLP binding were at sites in the ranges 57-90, 120-133, 140-151 and 255-256 (including the D77E and R149T mutations shown above to abrogate MHC-II mediated cell entry). However, several mutations at these sites in or near the core putative MHC-II binding pocket increased binding to tufted duck MHCII-VLPs, including D53C, I80L, E119G, and I121R. Other mutations that we measured to increase MHCII-VLP binding were located around sialic acid binding pocket (eg, sites 137, 155, and 187-227), raising the possibility that they may affect binding to other sialylated proteins on the MHCII-VLPs rather than the MHC-II protein itself.

**Figure 3.**
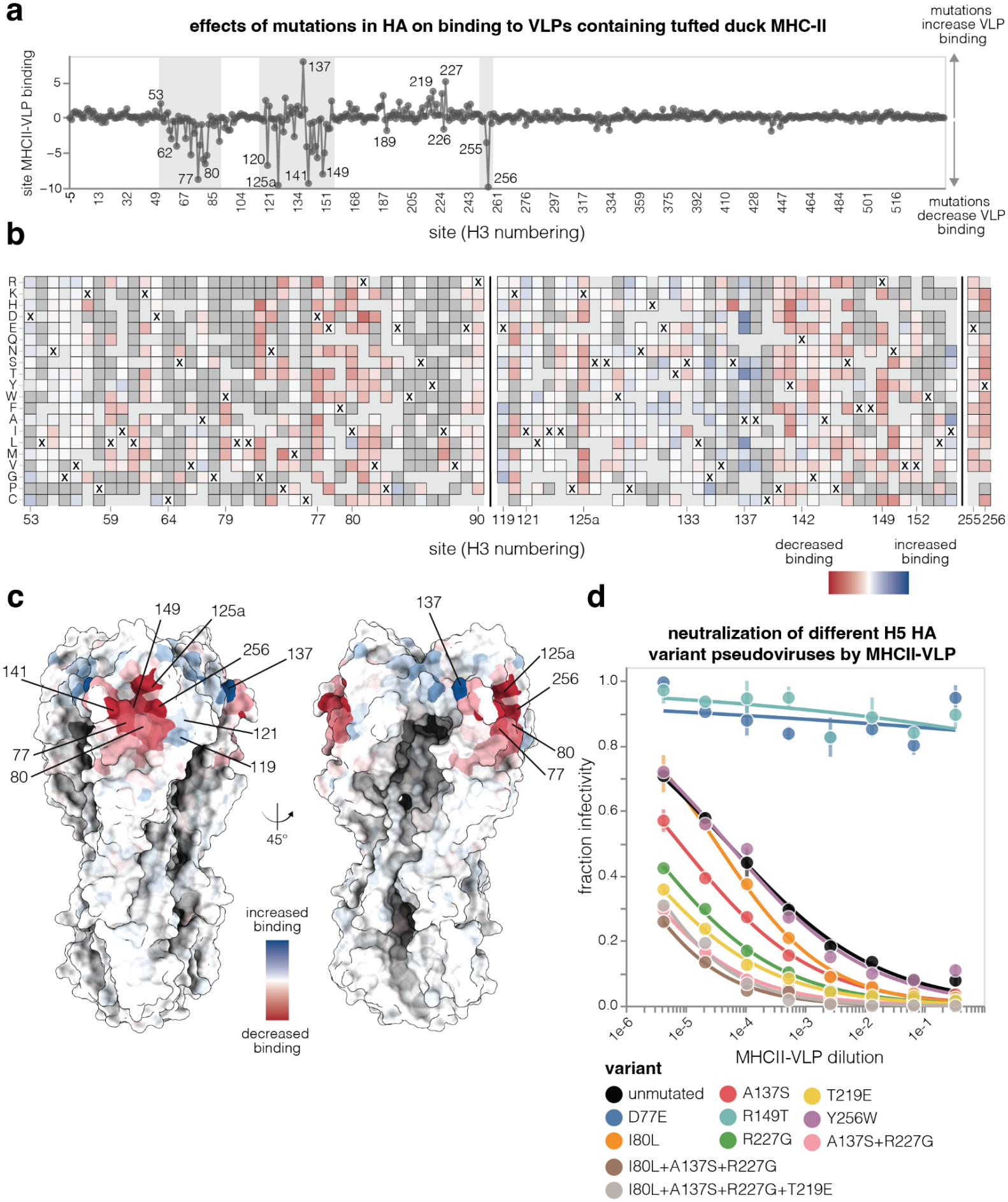
Effects of H5 HA mutations on binding to tufted duck MHCII-VLPs. **a**. Total effect of mutations at each HA site on tufted duck MHCII-VLP binding. Positive values indicate most mutations at a site increase MHCII-VLP binding, and negative values indicate most mutations at a site decrease MHCII-VLP binding. **b**. Effects of individual mutations on MHCII-VLP binding at key HA sites (indicated by gray shading in panel **a**). Negative values (red) indicate mutations decrease MHCII-VLP binding, positive values (blue) indicate mutations increase binding, dark grey indicates mutations too deleterious for pseudovirus entry in 293T cells to measure an effect on binding, and light grey indicates mutations for which no measurement was made. At each site, ‘x’ indicates the wildtype amino acid in the A/American Wigeon/South Carolina/USDA-000345-001/2021 H5 HA. See Supplementary Fig. 7c and the interactive plot at https://dms-vep.org/Flu-H5N1-American-Wigeon-2021-HA-tufted-duck-MHCII-DMS/binding.html for heatmaps covering the entirety of HA. **c**. Structure of H5 HA (PDB: 9DWE) colored by the total effect of all mutations at each site on MHCII-VLP binding. **d**. Validation assays showing neutralization of individually produced pseudoviruses with the indicated H5 HA mutants by MHCII-VLPs; HAs with reduced neutralization (eg, D77E, R149T) have reduced MHCII-VLP binding whereas HAs with increased neutralization (eg, I80L+A137S+R227G+T219E) have increased MHCII-VLP binding.

To validate the MHCII-VLP binding deep mutational scanning, we generated pseudoviruses carrying different HA mutations and measured their neutralization by MHCII-VLPs. Pseudoviruses with the D77E or R149T mutations in HA showed complete loss of neutralization by MHCII-VLPs, consistent with the expected strong decrease in binding to tufted duck MHC-II. Pseudoviruses with the I80L, A137S, R227G, or T219E mutations in HA showed the expected increase in MHCII-VLP neutralization. However, Y256W HA did not cause any measurable change in MHCII-VLP neutralization even though our deep mutational scanning suggests this mutation should decrease MHC-II binding.

### Effects of mutations to tufted duck MHC-II on HA binding

To define the sites in MHC-II that are critical for its interaction with HA, we first tested if the MHC-II / HA interaction is amenable to inverted pseudotyping^29,30^, an approach where lentiviral particles expressing the receptor protein infect cells that express the viral entry protein (**Supplementary Fig. 8a**). Indeed, pseudoviruses expressing tufted duck MHC-II (but no viral entry protein) could efficiently enter 293T cells expressing H5 HA but could not infect unmodified 293Ts (**Supplementary Fig. 8b**). This result confirms that the interaction between MHC-II and HA can mediate virion entry even in the inverted case when the MHC-II is expressed on the virion and the HA is expressed on the cell surface.

We leveraged this inverted pseudotyping to measure how mutations to tufted duck MHC-II affect its interaction with H5 HA by generating deep mutational scanning libraries^9^ where the pseudoviruses expressed different MHC-II mutants but no viral entry protein. The libraries were designed to have every possible single amino-acid mutation in the ectodomains of both the alpha and beta chains of tufted duck MHC-II, and the actual libraries covered almost all of these designed mutations (**Supplementary Fig. 8c**). To measure how mutations in MHC-II affect binding to HA, we incubated the MHC-II-expressing pseudovirus libraries with soluble H5 HA trimer under the expectation that pseudoviruses would be neutralized by soluble HA in proportion to the affinity of their mutant MHC-II for HA, analogous to the neutralization of the HA-expressing pseudoviruses by MHCII-VLPs described above. Indeed, soluble HA effectively neutralized the ability of MHC-II-expressing pseudoviruses to infect 293T-H5-HA cells, with MHC-II-expressing pseudoviruses being more potently neutralized by a mutant of HA that we showed in **Fig. 3d** to bind more strongly to MHC-II (**Supplementary Fig. 8d**).

The deep mutational scanning revealed that the mutations to MHC-II with the greatest effects on binding to the HA were all in the alpha chain, primarily in the α1 domain (**Fig. 4, Supplementary Fig. 9**, and interactive plots at https://dms-vep.org/Tufted-duck-MHCII-DMS/). Mutations at some of these alpha chain sites (eg, 15, 23, 69, and 74) increased MHC-II binding to HA, whereas mutations at other sites (eg, 22, 24, 37-46, and 57-68) decreased MHC-II binding. Mutations at several sites in the beta chain (eg, 15, 16, 53, 59, 78, and 87) also affected HA binding, but to a lesser extent than mutations in the alpha chain (**Fig. 4** and **Supplementary Fig. 9**). Note that MHC-II sites refer to sequential numbering of the tufted duck MHC-II ectodomain; these are not equivalent to homologous site numbers in human MHC-II.

**Figure 4.**
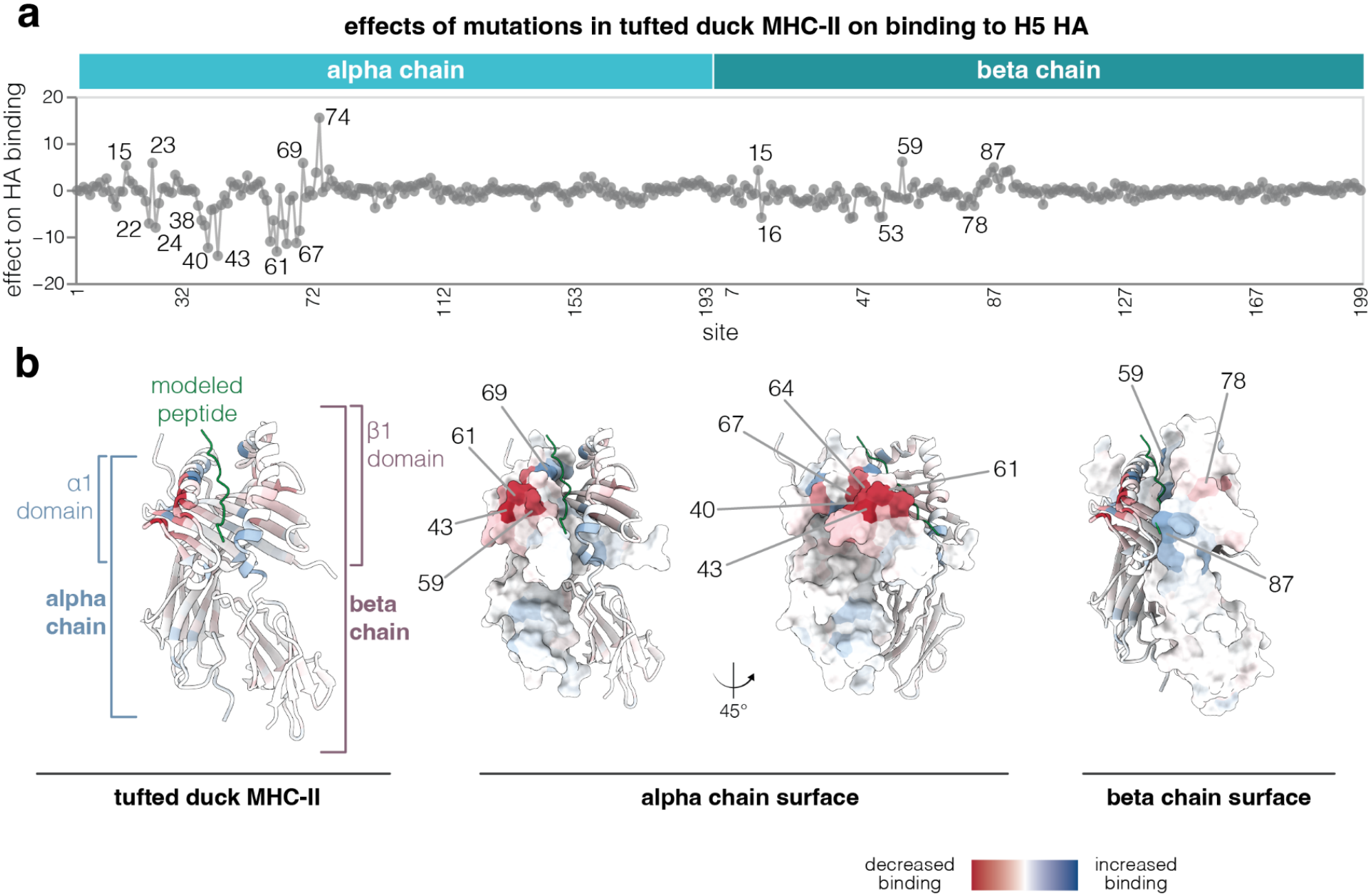
Effects of mutations in tufted duck MHC-II on H5 HA binding. **a**. Total effect of mutations at each site in the ectodomains of the alpha and beta chains of MHC-II on binding to H5 HA, as measured by pseudovirus deep mutational scanning. Sites are numbered starting with one for the first ectodomain site for both alpha and beta chains. Positive values indicate most mutations at a site increase HA binding, and negative values indicate most mutations at a site decrease HA binding. For an interactive version of this plot see https://dms-vep.org/Tufted-duck-MHCII-DMS/binding.html. **b**. AlphaFold 3-predicted structure of tufted duck MHC-II colored by the total effects of all mutations at each site on HA binding (red indicates most mutations at a site decrease HA binding). The peptide in green is aligned from the PDB:8JRJ structure and is shown here for visual orientation purposes.

We analyzed MHC-II alpha chains from several other avian and mammalian species for mutations at sites that the tufted duck MHC-II deep mutational scanning indicated were important for HA binding (**Supplementary Fig. 10a**). The MHC-II alpha chains from other species had mutations measured to both increase and decrease binding of tufted duck MHC-II to HA, with MHC-II from black swan, bald eagle, and human predicted to have modestly reduced H5 HA binding— but MHC-II from ferret, chicken, and mouse predicted to have strongly reduced H5 HA binding. Indeed, inverse pseudotyping of lentiviral particles with black swan, bald eagle, and human MHC-II yielded titers on 293T-H5-HA cells that were modestly lower than those for tufted duck MHC-II, whereas pseudotyping with ferret, chicken, and mouse MHC-II yielded very low titers (**Supplementary Fig. 10b-c**). These results suggest it may be possible to predict from the tufted duck MHC-II deep mutational scanning the ability of MHC-IIs from different species to interact with the particular H5 HA (A/American Wigeon/South Carolina/USDA-000345-001/2021) used in these experiments, although HAs from other influenza strains may interact differently with MHC-II from different species due to the genetic variation of HA at the MHC-II binding interface.

### Cryo-EM structure of MHC-II bound to H5 HA

We hypothesized that it might be easier to determine the structure of HA bound to MHC-II if we could make a higher affinity complex. Using the deep mutational scanning data, we designed an H5 HA variant with four mutations (I80L, A137S, T219E, and R227G) and a tufted duck MHC-II variant with three mutations (G23M, K42I, Q65F), all of which were expected to increase binding between the two proteins. HAs also had three mutations in the stalk (H355W, K380M, E432L) previously shown to stabilize the prefusion conformation^31,32^. We then produced soluble HA trimer and MHC-II protein with or without these mutations, and measured their affinities using bio-layer interferometry. The binding between HA and MHC-II was stronger when both proteins contained the putative affinity-enhancing mutations (**Supplementary Fig. 11a**). We also used negative-stain EM to examine a mix of soluble tufted duck MHC-II and soluble stabilized HA trimers with or without the affinity-enhancing mutations: in the 2D class averages, MHC-II binding occupancy was observed for 21% of HA particles when both proteins had the affinity-enhancing mutations, compared to only 3% of HA particles when both proteins lacked the affinity-enhancing mutations (**Supplementary Fig. 11b**).

We used cryo-EM to determine a 3.4-Å-resolution structure of the complex between H5 HA and tufted duck MHC-II with both proteins containing the affinity-enhancing mutations (**Fig. 5, Supplementary Fig. 11**). The cryo-EM structure shows that tufted duck MHC-II binds the side of the HA head, with the interaction mediated by the α1 domain of the MHC-II alpha chain (**Fig. 5b-c**). The MHC-II peptide-binding groove, which is formed by both the alpha and beta chains of MHC-II, faces HA. Three regions of the MHC-II alpha chain form the main contacts with the HA: beta-sheet residues 22–24, the loop formed by residues 37–44, and the peptide-groove-forming alpha helix spanning residues 59–74. Some density for a peptide bound in the groove near HA is apparent in the cryo-EM model, although the density is weak as expected if individual MHC-II molecules are bound to different peptides (**Supplementary Fig. 11f**).

**Figure 5.**
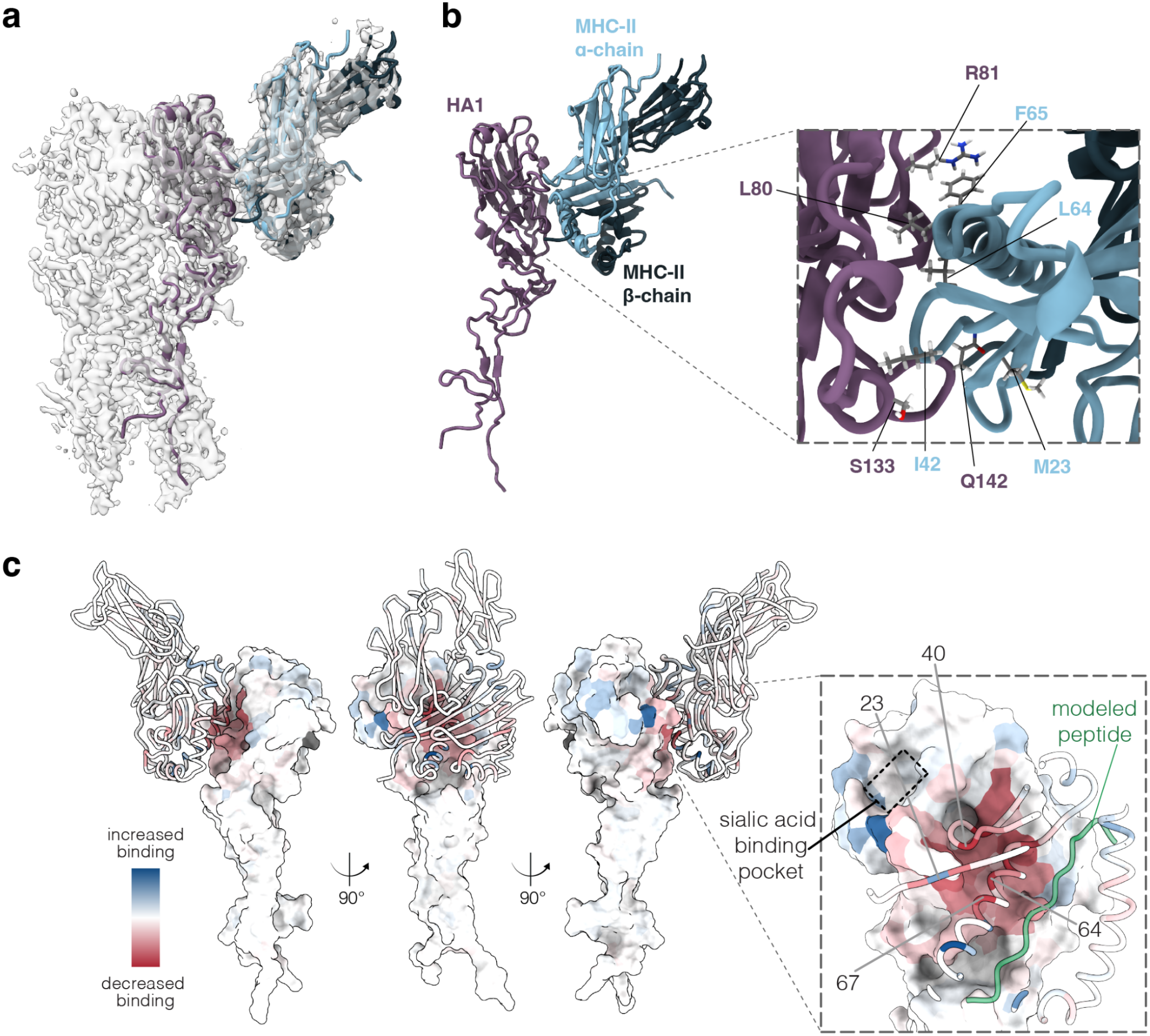
Cryo-EM structure of tufted duck MHC-II bound to H5 HA. **a**. 3.4 Å cryo-EM 3D reconstruction of HA in complex with MHC-II. The HA1 domain of HA is colored purple, MHC-II alpha chain is colored light blue and beta chain is colored dark blue. Note both proteins contain mutations identified in the deep mutational scanning as increasing the affinity of the complex. **b**. HA1 domain of HA bound to tufted duck MHC-II. The inset on the right shows a zoomed view of the core interaction surface. **c**. HA1 domain bound to tufted duck MHC-II, with both proteins colored by mutation effects on complex binding as measured by the deep mutational scanning. The inset on the right shows a zoomed-in view of the key interaction surface and includes a modeled peptide from PDB:1AQD for visual orientation purposes; peptide density is observed but cannot be modeled in the MHC-II structure presented in this manuscript.

The sites that our deep mutational scanning identified as important for MHC-II binding nearly all map to the interface between HA and MHC-II in the cryo-EM structure (**Fig. 5c**). HA residue R81 forms a cation-pi interaction with the mutated Q65F residue in MHC-II (**Fig. 5b-c**). HA mutation I80L promotes hydrophobic packing against L64 residue in MHC-II. The MHC-II mutation K42I promotes hydrophobic packing against the backbone of residue S133 in HA. Note however that several HA mutations that increased MHCII-VLP neutralization (eg, A137S, R227G, and T219E) are at sites that do not contact MHC-II in the structure, raising the possibility that they may have affected MHCII-VLP neutralization by interaction with sialic acid rather than MHC-II. Overall, the cryo-EM structure and deep mutational scanning data converge on the same architecture of the complex between HA and MHC-II—with the deep mutational scanning discovering the affinity-enhancing mutations that enabled determination of the cryo-EM structure.

### MHC-II interaction with HAs from H1, H2, H3, H7, and H9 subtypes

To examine if interaction of H5 HA with MHC-II extended to other influenza subtypes, we generated pseudoviruses bearing HAs from two primarily avian subtypes, H9 (A/quail/Hong Kong/G1/97) or H7 (A/Anhui/1/2013), and quantified their entry in no-sialic acid cells expressing human or tufted duck MHC-II (**Fig. 6a**). Pseudoviruses with either HA entered cells expressing tufted duck MHC-II far more efficiently than they entered control cells expressing no sialic acid (**Fig. 6a**). The pseudoviruses with H7 HA could also use human MHC-II for cell entry albeit much less efficiently than the tufted duck MHC-II, but the H9 HA pseudovirus could not utilize human MHC-II for cell entry.

**Figure 6.**
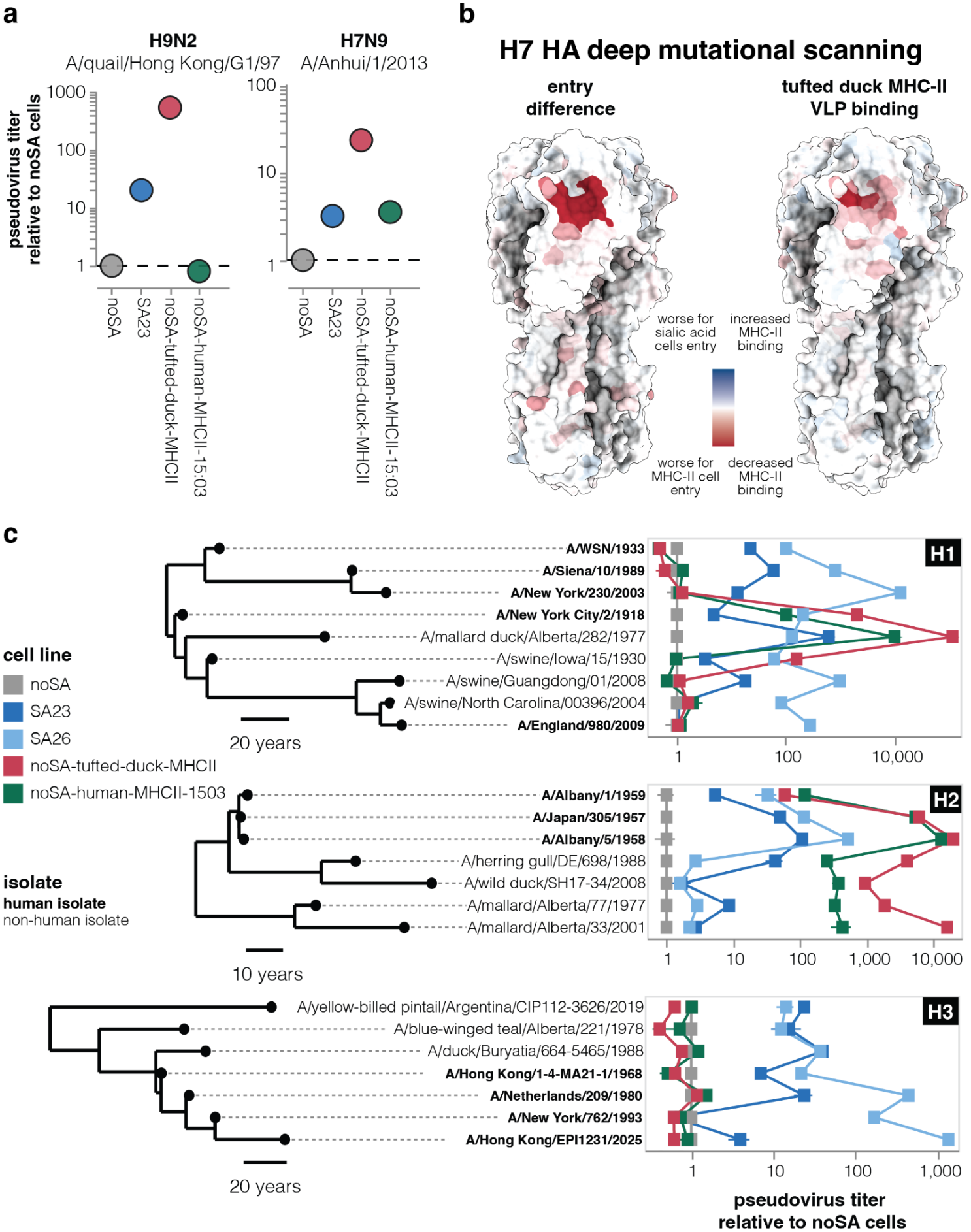
MHC-II-mediated cell entry by H9, H7, H1, H2, and H3 influenza HAs. **a**. Titers of lentiviral particles pseudotyped with the indicated H9 or H7 HA on cells expressing α2-3-linked sialic acids (SA23) or different MHC-II alleles relative to titers on no-sialic acid (noSA) cells. **b**. Structure of H7 HA (PDB: 6II9) colored by the effects of mutations on sialic acid versus tufted duck MHC-II mediated cell entry (left) or tufted duck MHCII-VLP binding (right) as measured by deep mutational scanning. In both cases red indicates sites where mutations are more deleterious for MHC-II mediated entry or binding, and blue indicates sites where mutations are more deleterious for sialic acid mediated entry or beneficial for MHC-II binding. See https://dms-vep.org/Flu-H7-Anhui13-MHCII-binding/ for the effects of individual H7 HA mutations on tufted duck MHC-II-mediated cell entry and binding. **c**. Titers of pseudoviruses with HAs from H1, H2 or H3 strains on no-sialic acid cells, cells expressing α2-3- or α2-6-linked sialic acids, or no-sialic acid cells expressing tufted duck or human MHC-II. Strains isolated from humans are indicated by bold letters. For the same plots showing raw titers rather than titers normalized to the noSA cells, see **Supplementary Fig. 12**. See https://dms-vep.org/Flu-H5N1-American-Wigeon-2021-HA-tufted-duck-MHCII-DMS/entry_titers.html for interactive versions of these titer plots.

We measured how nearly all mutations to the H7 HA affected tufted duck MHC-II mediated cell entry and binding to MHCII-VLPs using previously published H7 HA pseudovirus deep mutational scanning libraries^33^ and the same experimental workflow used for H5 HA earlier in this study. The sites in the H7 HA where mutations most strongly affected MHC-II-mediated cell entry or MHCII-VLP binding clustered just below the sialic acid–binding pocket in a region nearly identical to that identified by the H5 HA deep mutational scanning (**Fig. 6b** and interactive plots at https://dms-vep.org/Flu-H7-Anhui13-MHCII-binding/). This result indicates that the same HA region mediates MHC-II binding for both H5 and H7 HA subtypes.

We also examined MHC-II usage by HAs from the three influenza A subtypes known to have caused human pandemics followed by sustained endemic human transmission: H1, H2, and H3^34^. For each subtype we chose an early human pandemic strain, several later post-pandemic human strains, one or more avian strains, and in the case of H1 several swine strains (**Fig. 6c**). We then measured the ability of pseudoviruses expressing HA from each of these subtypes to enter no-sialic acid cells expressing MHC-II versus sialic acid expressing cells.

Among H1 HAs, the highest entry titers were on the tufted duck MHC-II expressing cells for pseudoviruses with HA from an avian strain, a human 1918 pandemic strain, and an early classical swine strain (**Fig. 6c**). In contrast, tufted duck MHC-II did not mediate efficient entry for pseudoviruses with HAs from later human H1N1 strains descended from the 1918 virus, later swine strains, or a 2009 human swine-origin H1N1 pandemic strain (**Fig. 6c**). Pseudoviruses with the 1918 HA could also use human MHC-II for cell entry, but with lower efficiency than tufted duck MHC-II.

Pseudoviruses with H2 HAs from avian and early pandemic (1957 and 1958) human strains could efficiently use both tufted duck and human MHC-II for cell entry, with the avian strains using tufted duck MHC-II with somewhat higher efficiency but the early human pandemic strains using tufted duck and human MHC-II with comparably high efficiency (**Fig. 6c**). These results are consistent with other recent work showing H2 influenza strains can enter cells via MHC-II^5^.

None of the human or avian H3 HAs we tested could mediate pseudovirus entry via tufted duck or human MHC-II, although a previous study has reported that some H3 strains can utilize human and swine MHC-II for cell entry^6^. Overall, these results suggest the patterns of MHC-II usage differ among HAs both between and within subtypes, and can change during the course of virus evolution in different hosts.

## Discussion

It has been known for several years that some influenza HAs can use MHC-II for cell entry^3–7^, but the molecular basis of this interaction has remained difficult to define using standard experimental approaches. Here we use a novel combination of deep mutational scanning and cryo-EM to characterize the molecular interface between an H5 HA and tufted duck MHC-II. Specifically, we used pseudovirus deep mutational scanning to measure how nearly all mutations to both HA and MHC-II affect their interaction. We then leveraged identification of mutations that increased binding between the two proteins to determine a 3.4 Å cryo-EM structure of the complex.

Together, the deep mutational scanning and cryo-EM structure show that a surface on the HA head distinct from the sialic acid-binding pocket engages MHC-II, with HA primarily interacting with residues in the MHC-II alpha chain. The fact that the core of the interaction involves the MHC-II alpha chain is important because while the MHC-II beta chain is among the most diverse proteins within and between jawed vertebrate species^14,15^, the alpha chain is generally less variable. However, our deep mutational scanning shows that mutations at some sites in the MHC-II beta chain proximal to the peptide-binding groove also affect the interaction with HA. The cryo-EM structure shows that the MHC-II peptide-binding groove faces HA in the protein complex, so one speculative idea is that the identity of the bound peptide could influence HA’s interaction with MHC-II, enabling variation in the beta chain to indirectly affect the interaction with HA by altering the repertoire of bound peptides.

We also used deep mutational scanning to show that an H7 HA interacts with MHC-II similarly to an H5 HA, confirmed prior reports that some H2 HAs can use MHC-II to enter cells^5^, and discovered that HAs from some H1 and H9 strains can also use MHC-II for cell entry. However, the ability of different HAs to use MHC-II from different species for cell entry varies even within a single viral subtype. For instance, most H5 HAs use tufted duck MHC-II better than the human MHC-II allele we tested, but a few H5 HAs have lost the ability to use both tufted duck and human MHC-II. Likewise, an avian H1 HA and the 1918 H1 HA use both tufted duck MHC-II and human MHC-II, while other H1 HAs do not use any MHC-II we tested. These strain-to-strain differences could arise as a simple byproduct of the fact that both HA and MHC-II are highly variable, with MHC-II under strong diversifying selection both within and between species due to its immune function^35–37^ and HA under strong selection for antibody escape^38,39^. However, it is also possible that HAs have adapted to interact with MHC-II alleles from specific host species. It is provocative to hypothesize that HA’s interactions with different MHC-II alleles could help shape viral tropism or immunogenicity in different species, or even among individuals within the same species harboring different MHC-II alleles.

However, the biological role of HA’s interaction with MHC-II remains unclear. Both prior work^3–7^ and our study have assayed the ability of MHC-II to serve as an alternative entry receptor. However, it should not be assumed that cell entry is the only biologically relevant function of HA’s interaction with MHC-II. With a few exceptions (such as bat and a few avian influenza viruses^3,4,7^), HAs that use MHC-II for cell entry can also use sialic acid as an entry receptor. Sialic acids are broadly expressed on the target cells relevant for influenza infection, whereas MHC-II is expressed primarily by professional antigen-presenting cells^40^ and some epithelial cells, with expression often increased under inflammatory conditions^5,41–43^. So while HA’s binding to MHC-II could further promote infection of certain cells, those cells should already be at least somewhat susceptible to viral infection due to their expression of sialic acid.

An alternative possibility is that the biological function of HA’s binding to MHC-II is something other than viral cell entry. For instance, the gp42 protein from Epstein–Barr virus binds to MHC-II as a co-receptor for B-cell entry, but there is also evidence that this binding can interfere with TCR recognition of peptide–MHC-II complex^44,45^. Although gp42 and HA bind distinct sites on MHC-II^45^, our structural model suggests that HA binding would likely sterically interfere with TCR engagement of MHC-II. Thus, HA’s binding to MHC-II could conceivably affect CD4 T-cell activation, which in turn could attenuate production of the neutralizing antibodies that are the main correlate of protection against influenza^46,47^. In addition, MHC-II binding could specifically promote infection of immune cells, affecting the immune response^48^. While further work is needed to determine the full biological role of the interaction between HA and MHC-II, our study takes a major step by defining the molecular interface and showing that MHC-II binding is widespread among HAs of several subtypes.

## Acknowledgments

We thank Henrik Clausen and Yoshiki Narimatsu from Copenhagen Center for Glycomics for sharing the noSA, SA23 and SA26 cell lines. This work was supported in part by the NIH/NIAID under 75N93021C00015 (to J.D.B.) and grant U19 AI181881 (to N.P.K. and J.D.B.). J.D.B. is an Investigator of the Howard Hughes Medical Institute. This work was also supported in part by Coefficient Giving (to N.P.K.). This research was supported by NIH P30 CA015704 of the Fred Hutch/University of Washington/Seattle Children’s Cancer Consortium, which includes the Genomics & Bioinformatics Shared Resource, RRID:SCR_022606. Sara Sunshine is a Washington Research Foundation postdoctoral fellow. Molecular graphics and analyses performed with UCSF ChimeraX, developed by the Resource for Biocomputing, Visualization, and Informatics at the University of California, San Francisco, with support from National Institutes of Health R01-GM129325 and the Office of Cyber Infrastructure and Computational Biology, National Institute of Allergy and Infectious Diseases.

## Supplementary Figures

**Supplementary Figure 1.**
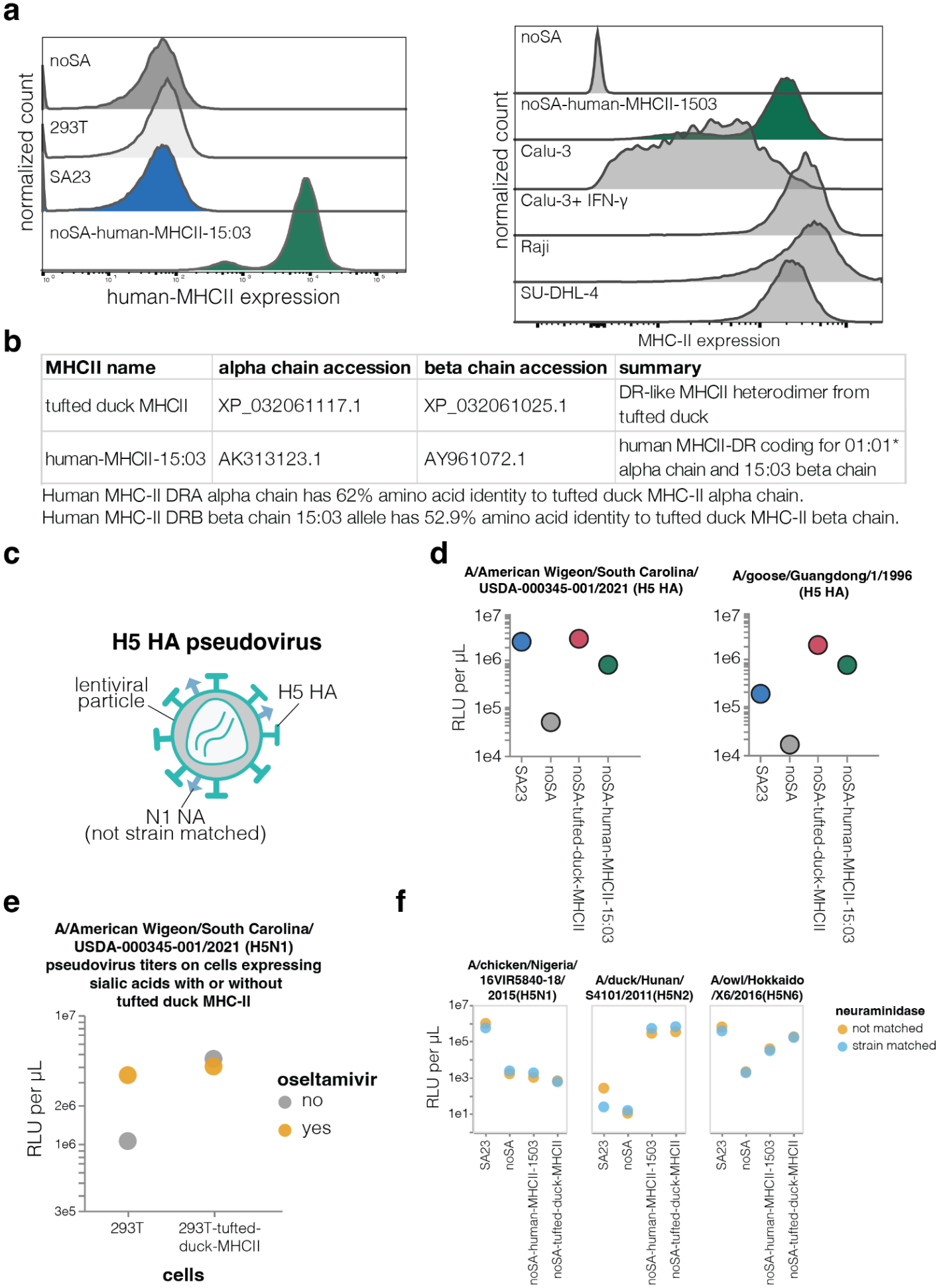
H5 HA pseudovirus entry in cells expressing different MHC-II alleles. **a**. Left, surface expression of human MHC-II in the cell lines used in this study (see **Fig. 1a**) quantified by flow cytometry using an antibody specific to the human MHC-II DRA alpha chain. Tufted duck MHC-II expression was not measured as no antibody that binds to this MHC-II is available. Right, comparison of MHC-II surface expression in the cell lines used in this study versus human B-cell lines (Raji and SU-DHL-4) or the Calu-3 human airway epithelial cell line with or without IFN-γ treatment. **b**. Details on the MHC-II alleles used in this study. **c**. A schematic of H5 HA pseudovirus. Lentiviral particles are pseudotyped with HA from a given H5 strain and the same N1 NA from A/turkey/Indiana/22-003707-003/2022 strain is used for all pseudoviruses. **d**. The same pseudovirus entry data as in **Fig. 1b** but showing raw titers in relative light units (RLU) per µl rather than titers normalized to the noSA cells. The low but measurable titers in the noSA could be because these cells may still express some residual sialic acid due to expression of alternative sialyltransferases^12^. **e**. Titers of lentiviral particles pseudotyped with A/American Wigeon/South Carolina/USDA-000345-001/2021 measured in unmodified 293T cells (which express sialic acid) and 293T cells overexpressing tufted duck MHC-II in the presence or absence of neuraminidase inhibitor oseltamivir. **f**. Titers of pseudoviruses made with strain-matched NA or NA from the A/turkey/Indiana/22-003707-003/2022 (H5N1) strain on different cell lines.

**Supplementary Figure 2.**
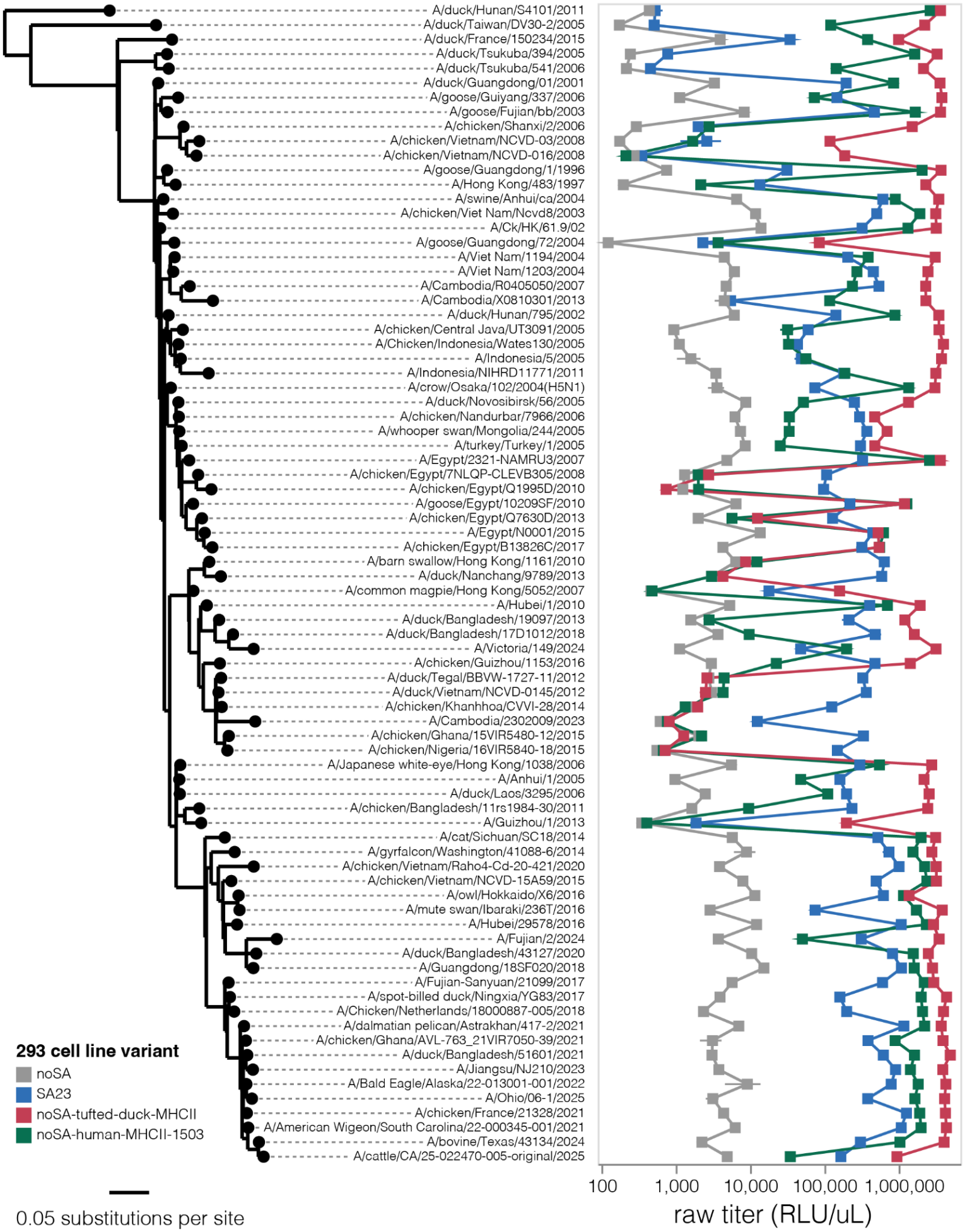
Non-normalized titers of pseudoviruses expressing H5 HAs on each cell line. The same data as in **Fig. 1c** but showing the raw titers in relative light units (RLU) per µl rather than titers normalized to the noSA cells. To view this plot interactively see https://dms-vep.org/Flu-H5N1-American-Wigeon-2021-HA-tufted-duck-MHCII-DMS/entry_titers.html.

**Supplementary Figure 3.**
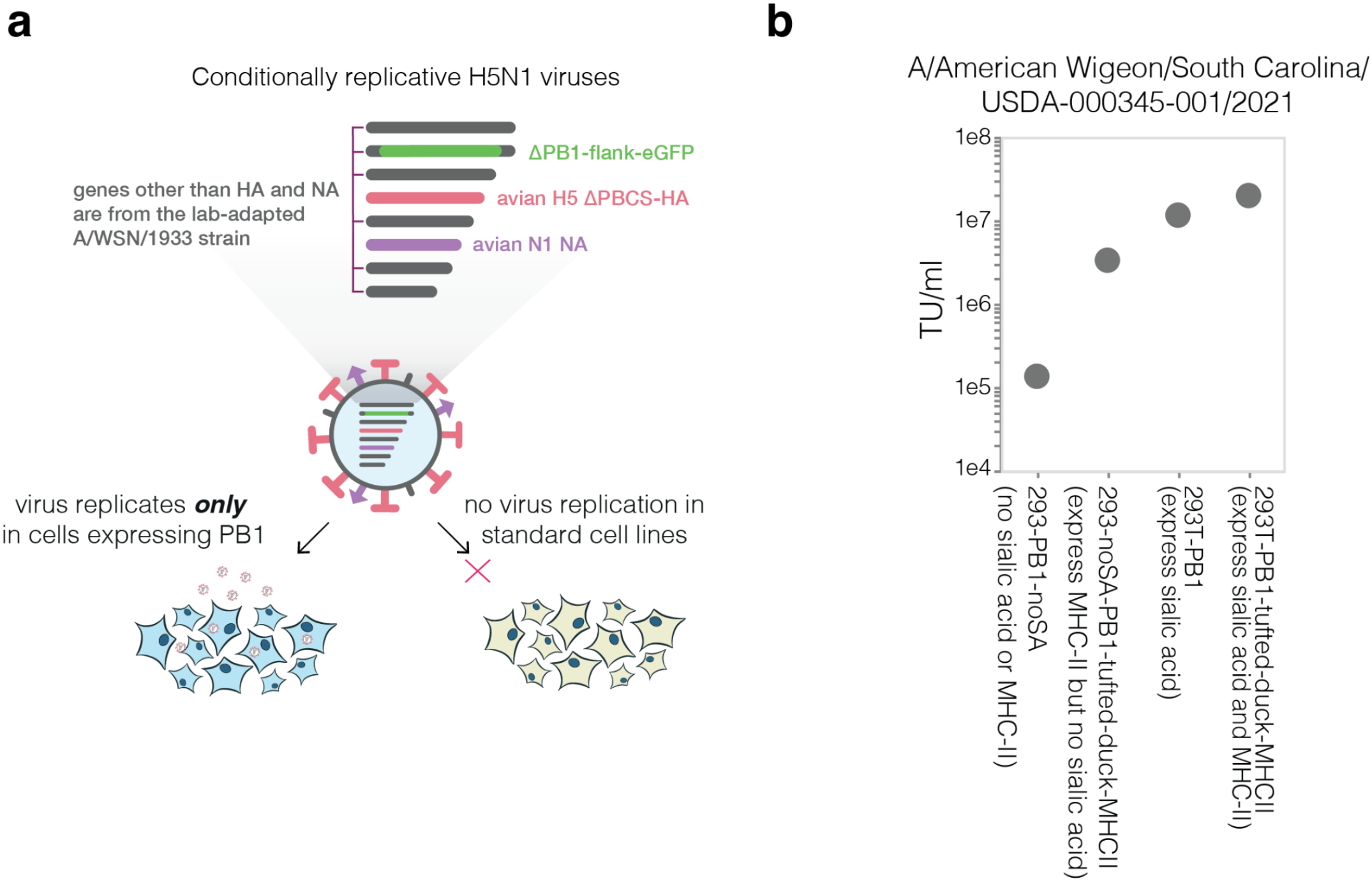
Entry titers of conditionally replicative influenza virus on cells expressing tufted duck MHC-II. **a**. Conditionally replicative influenza viruses encode avian H5 HA and N1 NA, five of the six remaining genomic segments from the lab-adapted A/WSN/1933 strain, and GFP in place of the PB1 polymerase gene^22^. These virions can only replicate in cells engineered to constitutively express the PB1 protein missing from the viral genome. As an additional biosafety precaution, the polybasic cleavage site is removed from the HA segment. **b**. Entry titers for a single round infection of the indicated cells with conditionally replicative influenza virus carrying HA from A/American Wigeon/South Carolina/USDA-000345-001/2021 (H5N1) strain. All cell lines express PB1. The 293-noSA-PB1 cells express no sialic acids and no MHC-II. The 293-noSA-PB1-tufted-duck-MHCII cells express no sialic acid but are engineered to constitutively express tufted duck MHC-II. 293T-PB1 cells express sialic acid but no MHC-II. 293T-PB1-tufted-duck-MHCII cells express sialic acids and tufted duck MHC-II. Titers were quantified as transcriptionally active virions per µl as quantified by expression of the GFP encoded in place of PB1.

**Supplementary Figure 4.**
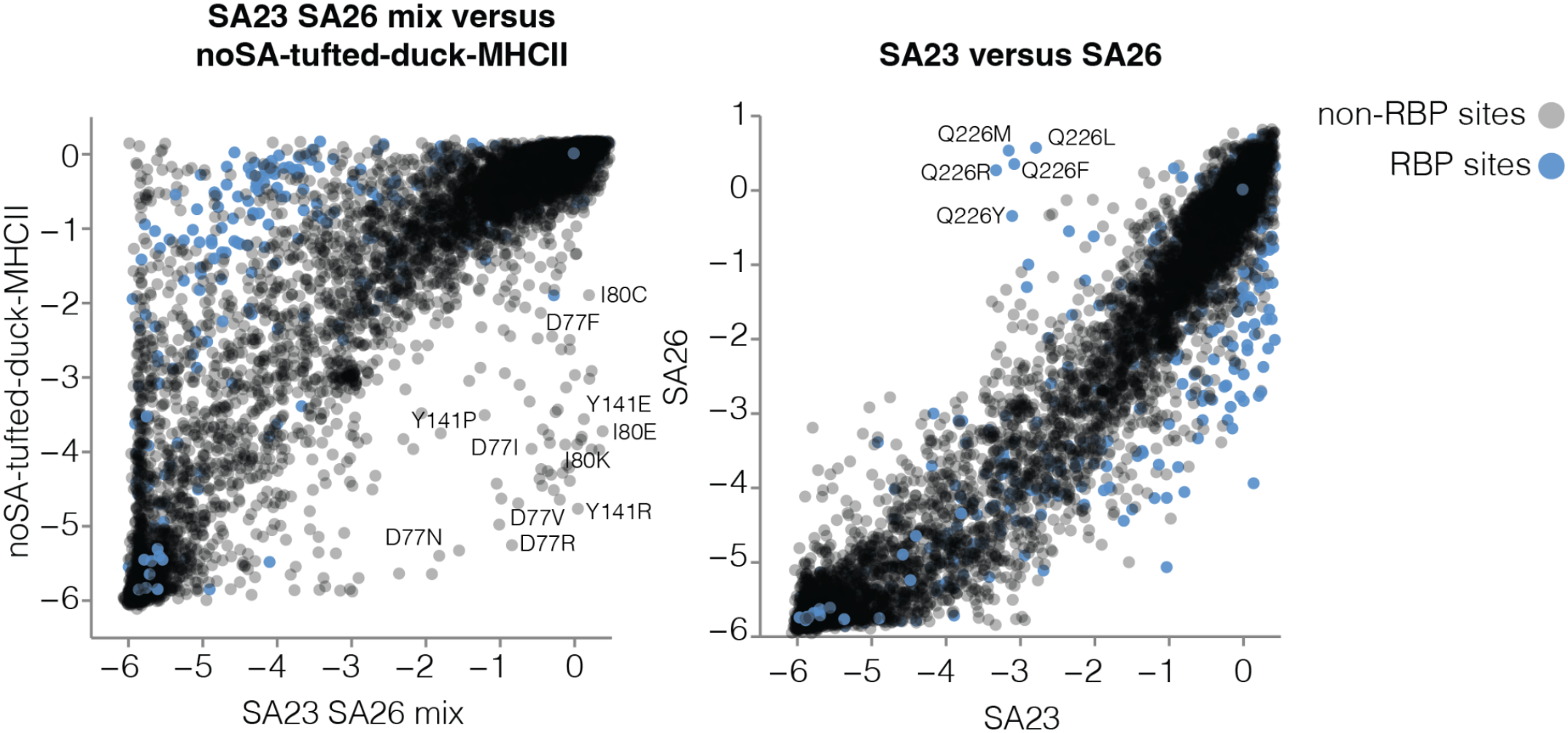
Effects of H5 HA mutations on pseudovirus entry in different cells. Scatter plots showing effects of mutations on entry in noSA-tufted-duck-MHCII cells versus a mix of SA23 and SA26 cells (left panel), or SA23 cells versus SA26 cells (right panel), as measured by pseudovirus deep mutational scanning. A negative effect indicates a mutation impairs entry in that cell. Sites that form the sialic acid receptor binding pocket (RBP) are colored blue. See https://dms-vep.org/Flu-H5N1-American-Wigeon-2021-HA-tufted-duck-MHCII-DMS/entry.html for an interactive version of this plot that allows mousing over points for mutation details.

**Supplementary Figure 5.**
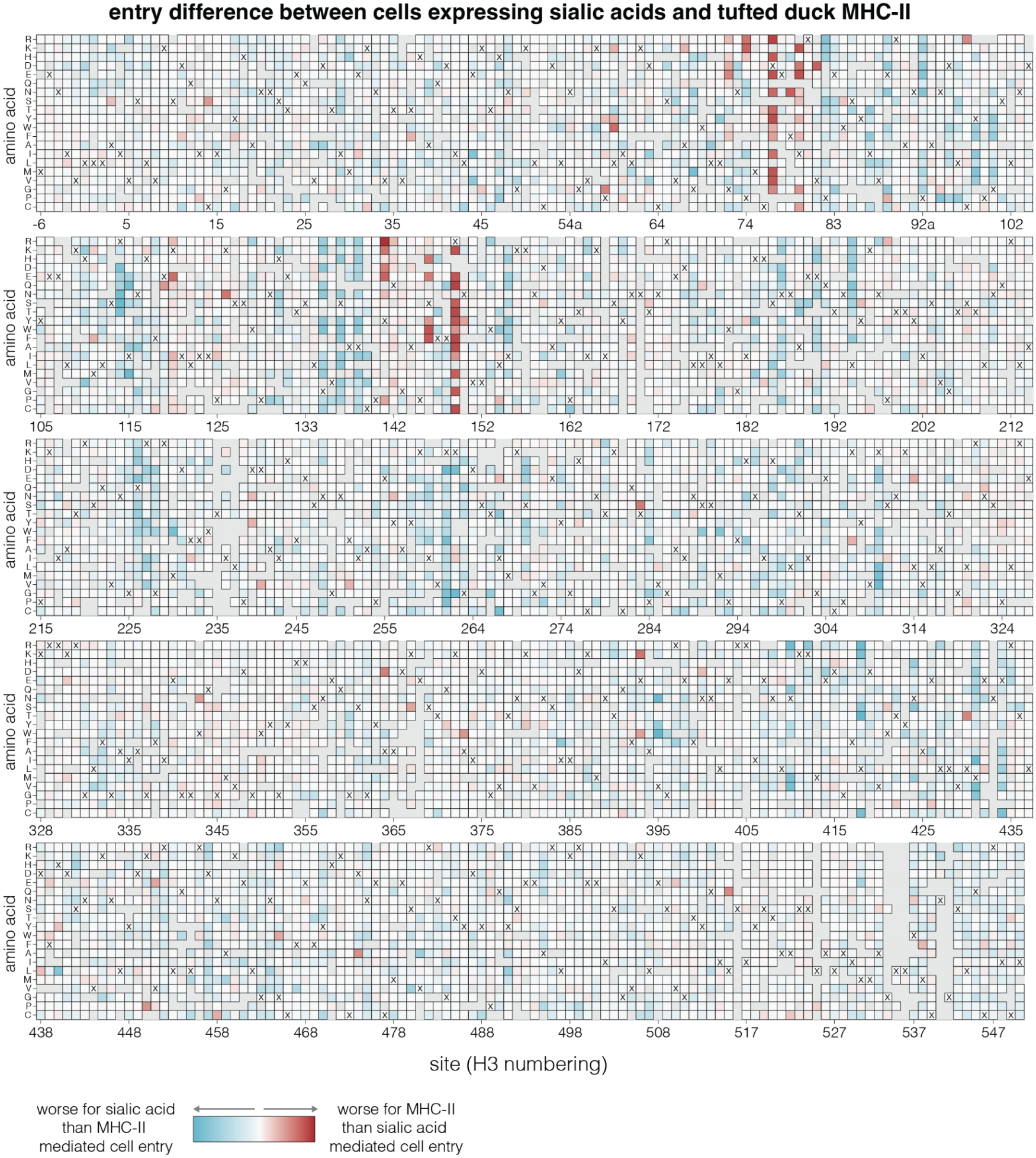
Difference between H5 HA mutation effects on entry in cells expressing sialic acids versus tufted duck MHC-II. Difference in mutation effects on pseudovirus entry in noSA-tufted-duck-MHCII cells versus a mix of SA26 and SA23 cells. Red indicates mutations that are more deleterious for entry in tufted duck MHC-II expressing cells than sialic acid expressing cells, and blue indicates mutations that are more deleterious for entry in sialic acid expressing cells than MHC-II expressing cells. Gray indicates the effect of that mutation was not measured in our deep mutational scanning, and ‘x’ indicates wild type amino acid in the A/American Wigeon/South Carolina/USDA-000345-001/2021 HA. See https://dms-vep.org/Flu-H5N1-American-Wigeon-2021-HA-tufted-duck-MHCII-DMS/entry.html for an interactive version of this plot and heatmaps that show the absolute effects of mutations on entry in each type of cell.

**Supplementary Figure 6.**
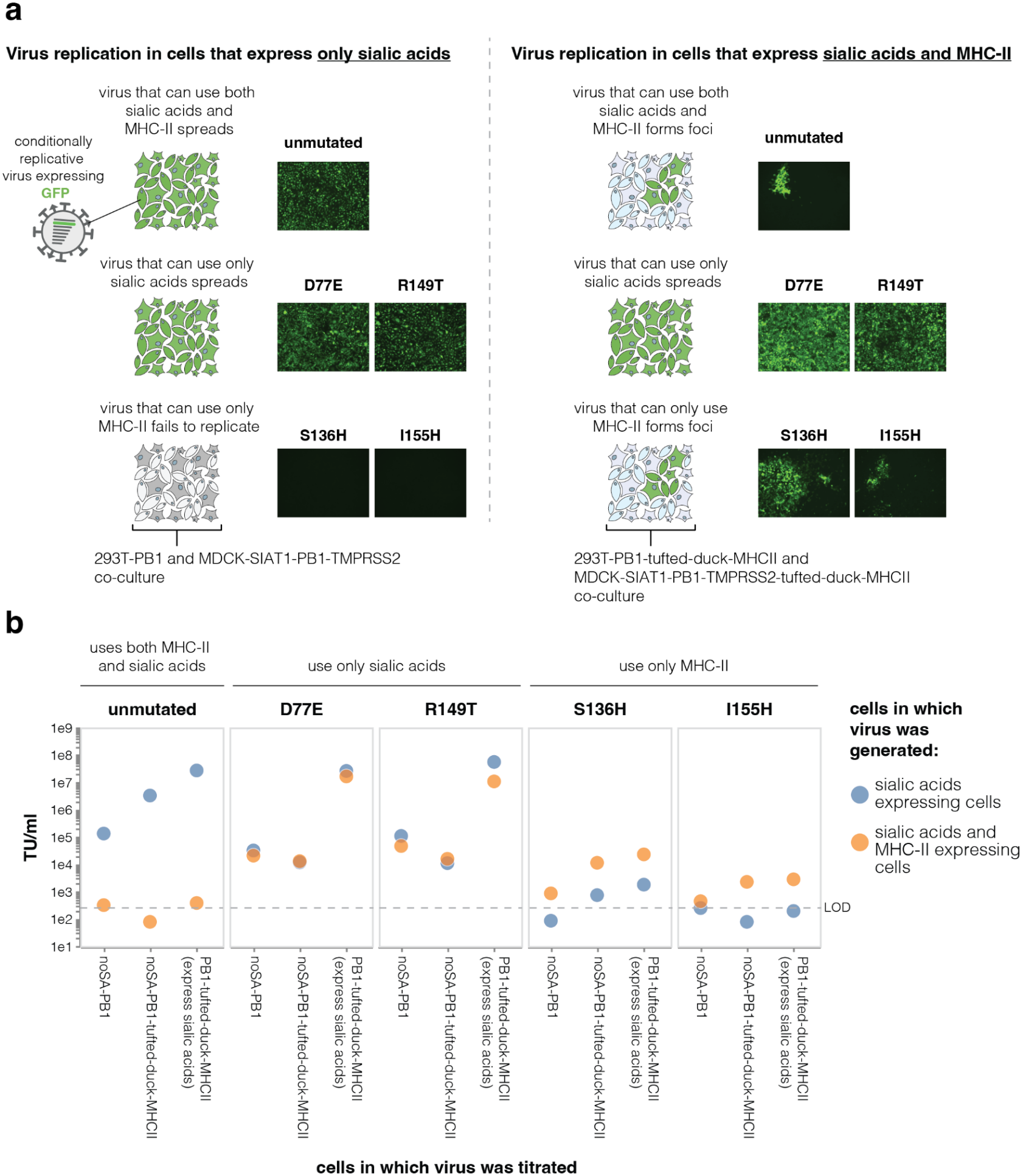
Titers of conditionally replicative influenza virus with HA mutations that ablate MHC-II or sialic acid mediated cell entrya. Multi-cycle replication of conditionally replicative influenza viruses expressing GFP in place of PB1^22^ (see **Supplementary Fig. 3a**) on cells that express only sialic acid or cells that express both sialic acid and tufted duck MHC-II. When virus is produced and grown in cells that express only sialic acid, viruses with HAs that can use both sialic acid and MHC-II (unmutated) or only sialic acid (D77E and R149T) spread rapidly as visualized by the GFP expressed from the viral genome, whereas viruses with HAs that cannot use sialic acid (S136H and I155H) do not grow in sialic acid-only expressing cells. When virus is produced and grown in cells that express both sialic acid and tufted duck MHC-II, viruses with HAs that can use both MHC-II and sialic acid (unmutated) or only MHC-II (S136H and I155H) form localized foci presumably because the virions enter the cells but are not efficiently released due to binding of virion HA to cell-surface MHC-II. In contrast, viruses with HAs that only bind to sialic acids (D77E and R149T) can efficiently spread in cells expressing both sialic acid and MHC-II. **b**. Titers of viruses produced in different cell lines during the multi-cycle infection shown in **a** measured in a single-cycle infection on different cell lines. The noSA-PB1 cells express neither sialic acid nor MHC-II, the noSA-PB1-tufted-duck-MHCII cells express only MHC-II, and the PB1-tufted-duck-MHCII cells express both sialic acid and MHC-II. Titers were quantified as transcriptionally active virions per ul as quantified by expression of the GFP in target cells by flow cytometry. The dashed line indicates the limit of detection (LOD) for virus titers.

**Supplementary Figure 7.**
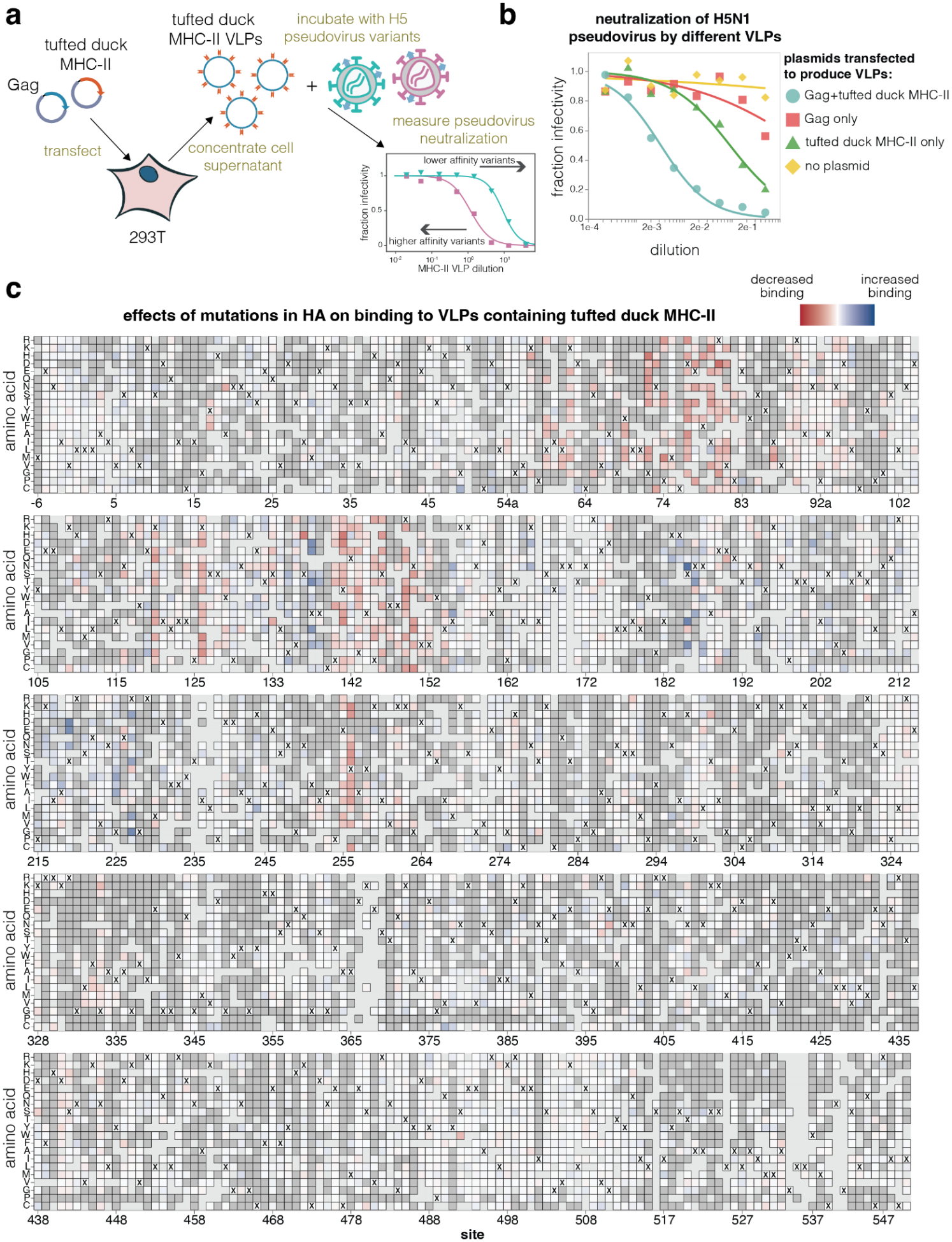
Tufted duck MHCII-VLP neutralization to measure effects of HA mutations on MHC-II bindinga. Tufted duck MHCII-VLPs were generated by transfecting 293T cells with expression plasmids for lentiviral Gag protein and tufted duck MHCII. After two days, the cell supernatant was collected and concentrated to collect tufted duck MHCII-VLPs. MHCII-VLPs were used to perform neutralization assays by incubating H5 pseudovirus variants with serial dilutions of MHCIII-VLPs and measuring changes in virus infectivity on 293T cells. **b**. Neutralization of H5 HA pseudovirus with cell supernatant from different VLP production conditions or controls. Only MHCII-VLPs produced by cotransfection of tufted duck MHC-II and Gag were potent pseudovirus neutralizers, although some neutralization was observed by supernatant of cells transfected with tufted duck MHC-II alone, likely due to some MHC-II shedding from transfected cells. **c**. Effects of all measured HA mutations on MHCII-VLP binding. This panel is similar to **Fig. 3b** but shows the full HA. See https://dms-vep.org/Flu-H5N1-American-Wigeon-2021-HA-tufted-duck-MHCII-DMS/binding.html for an interactive version of this heatmap.

**Supplementary Figure 8.**
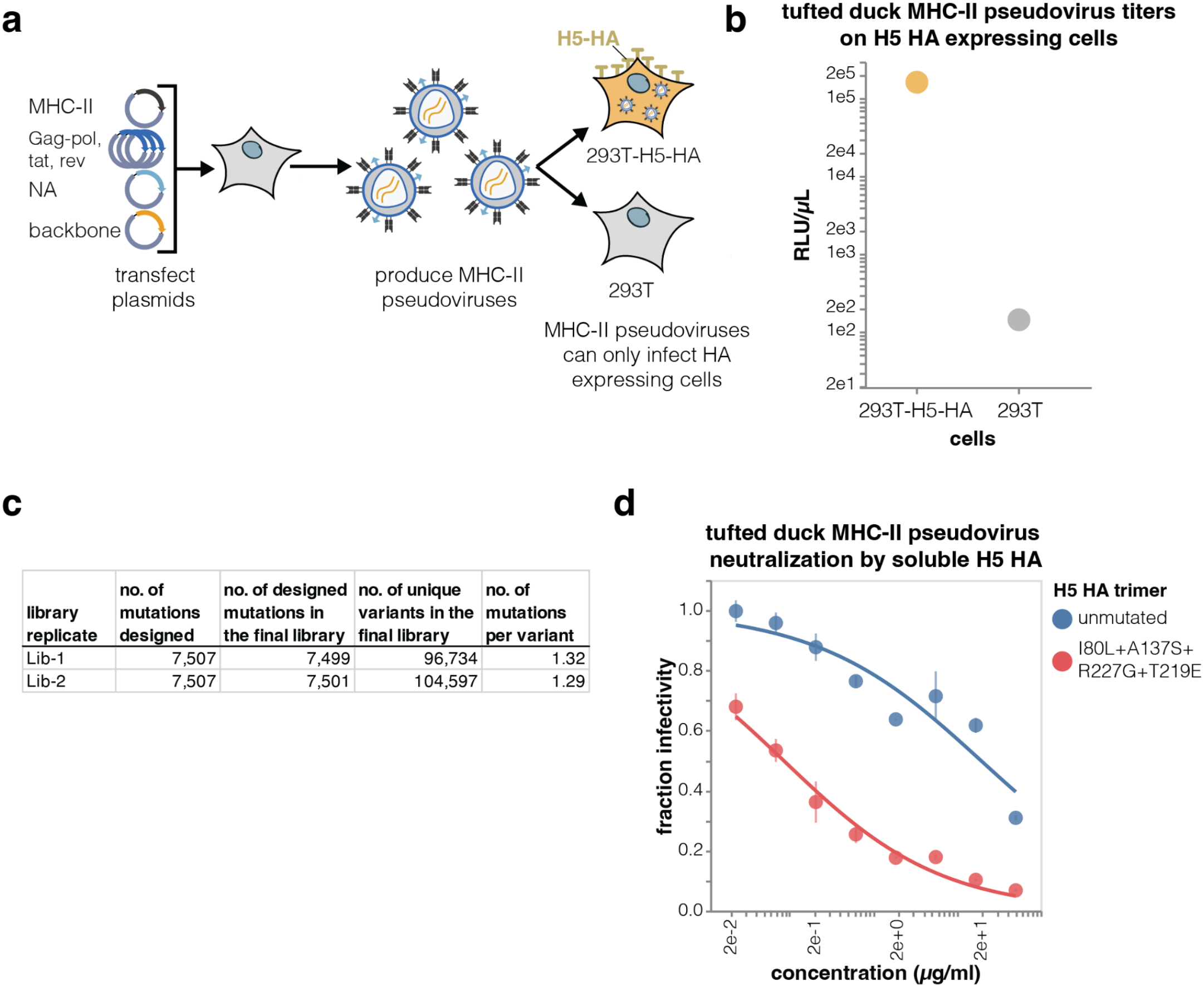
MHC-II inverted pseudotyping and deep mutational scanninga. Inverted pseudotyping of lentiviral particles with tufted duck MHC-II. 293T cells were transfected with plasmids encoding lentiviral helper proteins, a lentiviral backbone, and expression plasmids coding for MHC-II and influenza neuraminidase (NA). Pseudovirus-containing cell supernatant was collected 48 hours later and titrated on 293T cells that do or do not express H5 HA from A/American Wigeon/South Carolina/USDA-000345-001/2021 strain. The MHC-II pseudoviruses can only enter cells that express HA. Note that neuraminidase (NA) is co-transfected during lentivirus production to reduce sialic acid mediated entry in HA expressing cells (see Methods for details). **b**. Tufted duck MHC-II pseudovirus titers on 293T and 293T-H5-HA cells. **c**. Composition of the two independent tufted duck MHC-II deep mutational scanning libraries. The table shows the total number of designed ectodomain mutations, the number of designed MHC-II mutations successfully introduced in each library, the number of unique barcoded library variants, and the per variant mutation rate. Note that the number of barcoded variants exceeds the number of unique mutations because most mutations are associated with multiple barcoded variants. **d**. Neutralization of tufted duck MHC-II pseudovirus infection into 293T-H5-HA by soluble trimers of unmutated H5 HA trimer or a variant of H5 HA with mutations shown earlier in this study to increase binding to tufted duck MHC-II. As expected, soluble HA that binds more strongly to MHC-II more potently inhibits infection by the MHC-II pseudoviruses.

**Supplementary Figure 9.**
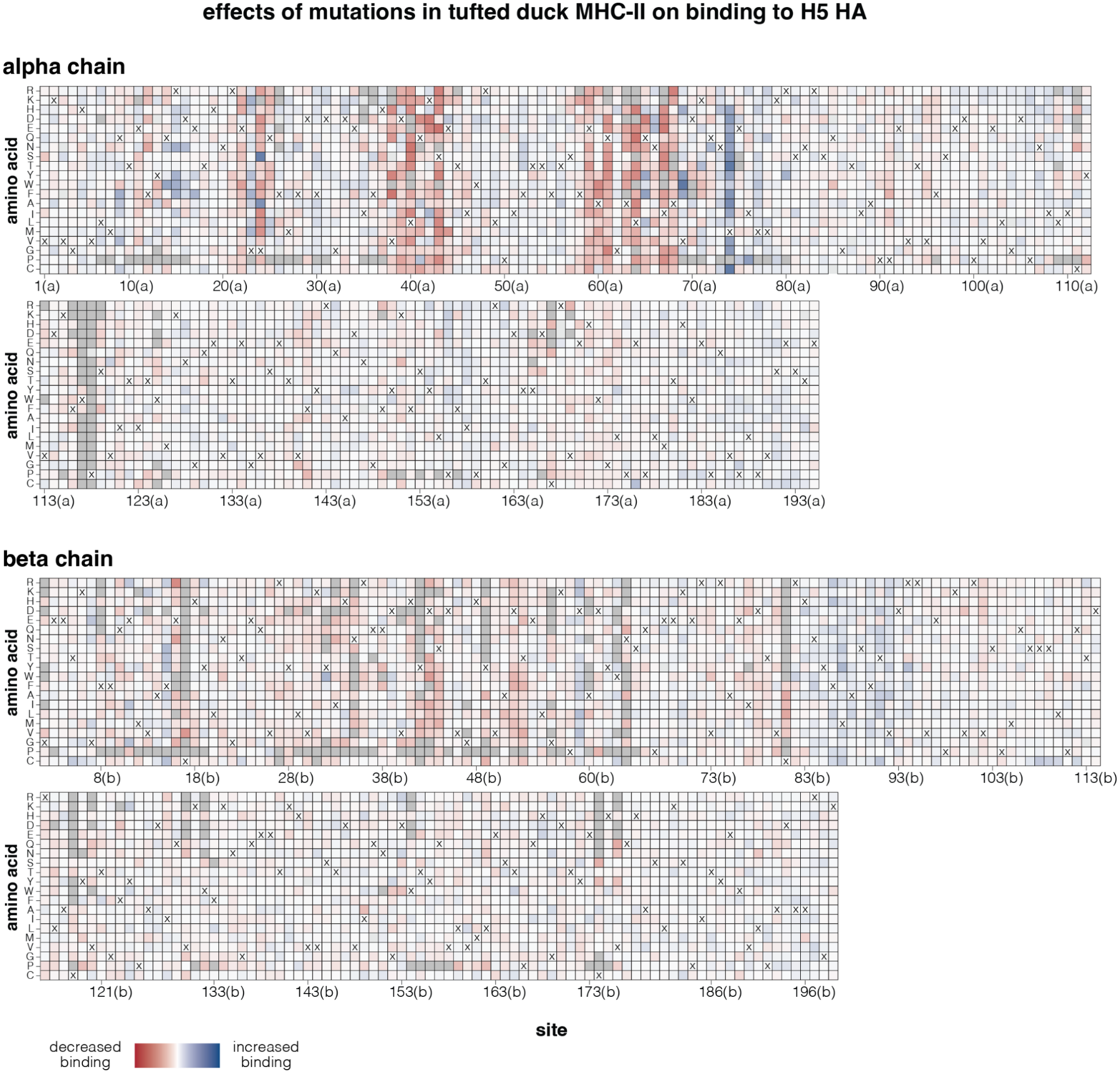
Effects of mutations in tufted duck MHC-II on binding to H5 HA. Effects of MHC-II mutations on H5 HA binding as measured by the inverted pseudovirus deep mutational scanning. Negative values (red) indicate mutations that decrease HA binding, positive values (blue) indicate mutations that increase binding, dark grey indicates MHC-II mutations that strongly impair pseudovirus entry in 293T-H5-HA cells (likely by causing MHC-II misfolding), and light grey indicates mutations for which no measurement was made (there are four such mutations: E85C, I25F, V143Y, Q150M). At each site, ‘x’ indicates the wildtype amino acid in tufted duck MHC-II. See https://dms-vep.org/Tufted-duck-MHCII-DMS/binding.html for an interactive version of this plot.

**Supplementary Figure 10.**
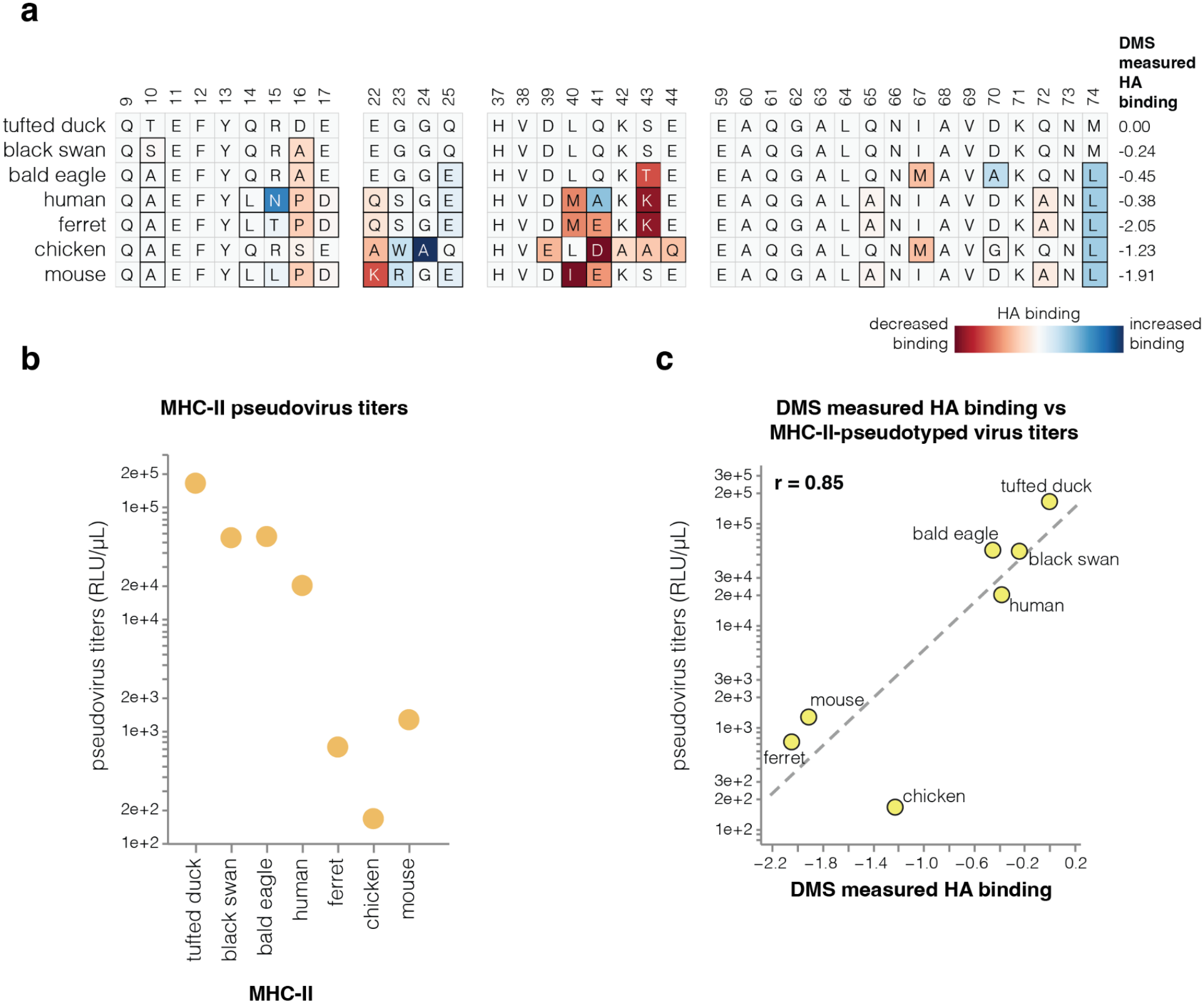
Interaction of MHC-IIs from different species with H5 HAa. Sequence alignment of key MHC-II alpha chain sites involved in the interaction with HA as determined by the inverted pseudovirus deep mutational scanning. Mutations relative to the tufted duck MHC-II sequence are colored based on their effect on H5 HA binding as measured by deep mutational scanning. The sum of the deep mutational scanning (DMS) measured effects of mutations at these sites is shown to the right of the alignment. **b**. Titers of lentiviral particles pseudotyped with MHC-IIs from different species on 293T-H5-HA cells. **c**. Relationship between the titers of lentiviral particles pseudotyped with MHC-IIs from different species measured on 293T-H5-HA cells and deep mutational scanning measured HA binding scores (computed by summing the effects of all mutations in that MHC-II relative to the tufted duck MHC-II in the region indicated in panel **a**). r indicates Pearson correlation, dashed line linear regression.

**Supplementary Figure 11.**
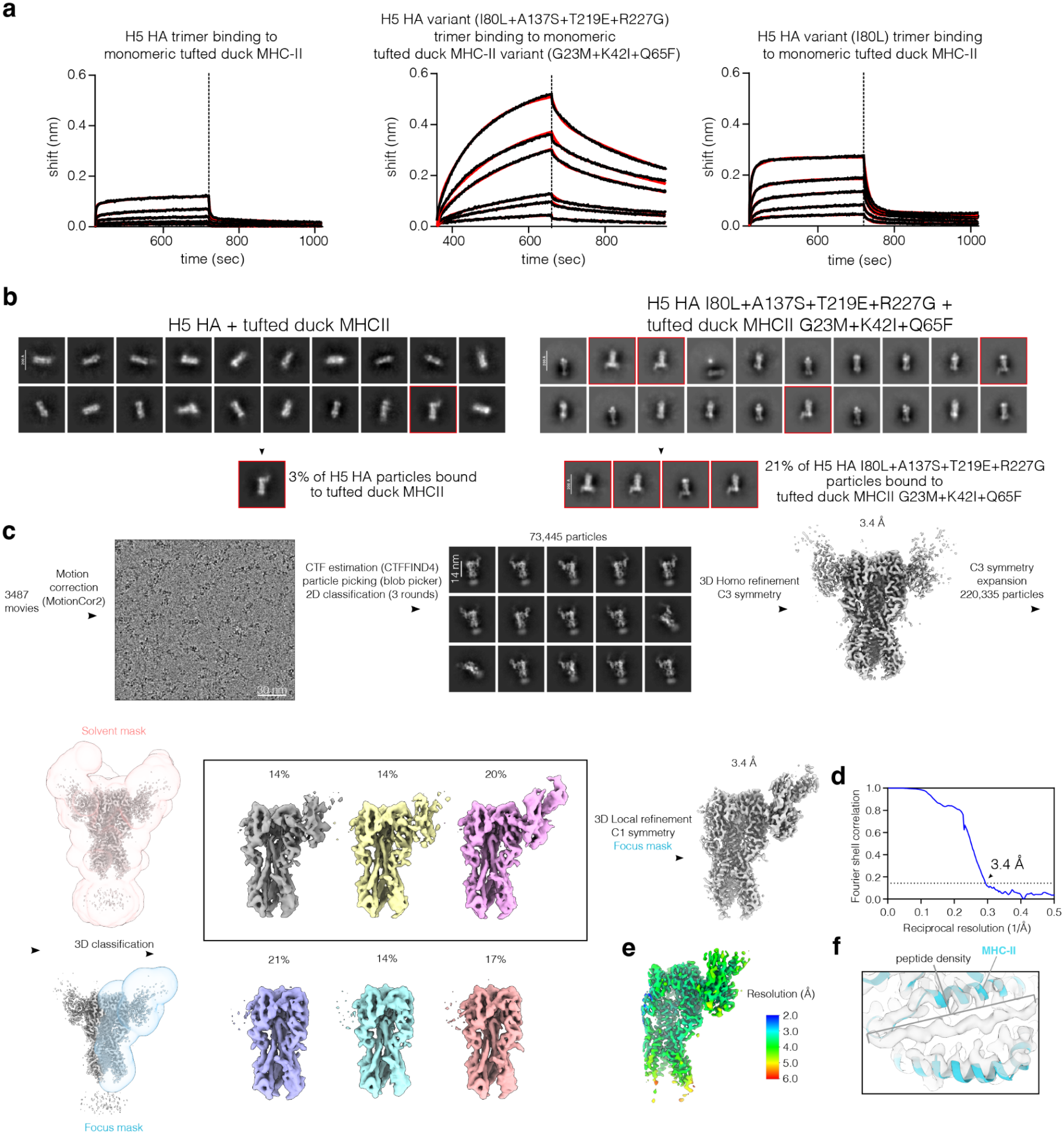
Measurements for H5 HA binding to tufted duck MHC-II, negative stain EM 2D class averages and cryo-EM processing workflow for complex of H5 HA bound to tufted duck MHC-II. **a**. Bio-Layer interferometry (BLI) of tufted duck MHC-II binding to immobilized H5 HA trimers without (left) or with (right) mutations in HA and MHC-II that increase the binding affinity. The BLI sensorgrams (black lines) were fit (red lines) using a 2:1 binding model. A 2:1 model was necessary to achieve a good fit. Measured affinities were K_D1_=3.5 µM and K_D2_=0.6 µM for unmutated HA binding to unmutated MHC-II, K_D1_=1.3 µM and K_D2_=2 µM for variant HA (I80L, A137S, T219E, R227G) binding to variant MHC-II (G23M, K42I, Q65F), and K_D1_=1.04 µM and K_D2_=0.1 µM for variant HA (I80L) binding to unmutated MHC-II. The two KDs may reflect specific HA–MHC-II binding and nonspecific interactions between MHC-II and the sensors, as high concentrations of MHC-II were required for this assay. **b**. Negative stain EM 2D class averages of HA trimers without (left) or with (right) the four affinity-enhancing mutations (I80L, A137S, T219E, R227G) incubated with tufted duck MHC-II with or without the three affinity-enhancing mutations (G23M, K42I, Q65F). Red boxes indicate class averages with tufted duck MHC-II bound. **c**. Cryo-EM processing diagram, **d**. gold-standard FSC curve, and **e**. Local resolution estimation for the reconstruction of the complex of H5 HA with the four affinity-enhancing mutations in complex with tufted duck MHC-II with three affinity-enhancing mutations. In **d**, the arrow points to the reciprocal resolution where the FSC is at the 0.143 cutoff value, and the displayed value is the calculated resolution at that cutoff. **f**. Peptide density observed in MHC-II in cryo-EM reconstruction.

**Supplementary Figure 12.**
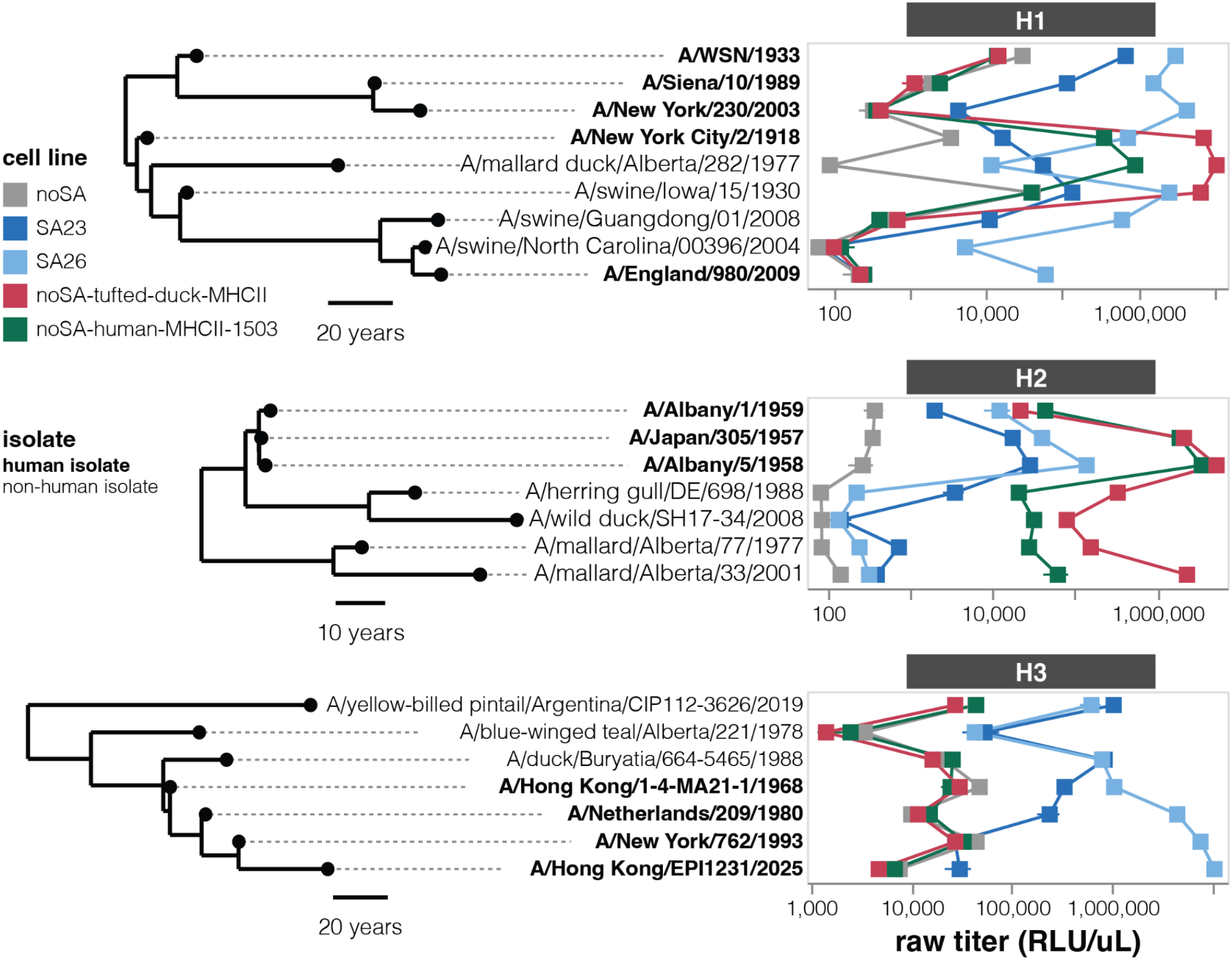
Non-normalized titers of pseudoviruses expressing H1, H2 and H3 HAs on each cell line. The same data as in **Fig. 6c** but showing the raw titers in relative light units (RLU) per µl rather than titers normalized to the noSA cells. See https://dms-vep.org/Flu-H5N1-American-Wigeon-2021-HA-tufted-duck-MHCII-DMS/entry_titers.html for interactive versions of these plots.

## Methods

### Biosafety and biosecurity statement

Most experiments in this study used lentiviral particles pseudotyped with influenza HA and neuraminidase (pseudoviruses). These pseudoviruses encode no viral proteins other than HA, and so can only undergo a single round of cell entry—meaning they provide a safe way to study the effects of mutations to viral proteins outside the context of an actual pathogen^49,50^. A few loss-of-function mutations for sialic acid or MHC-II-mediated entry identified using the pseudoviruses were validated in conditionally replicative influenza virions^22^; as detailed in **Supplementary Fig. 3a** these virions lack the essential viral PB1 polymerase protein and so can only replicate in a lab-engineered cell line that expresses PB1; in addition the non-HA/NA genes in these virions are from the lab-adapted A/WSN/1933 strain. Since neither the pseudoviruses nor conditionally replicative influenza virions can replicate in humans or any other animal, the experiments were performed at biosafety-level-2 in accordance with a Fred Hutch approved biosafety protocol.

We recognize concerns that even safe experiments could pose a so-called “information hazard” by providing knowledge that could be misused by others^51,52^. Because it remains unclear if or how MHC-II binding contributes to the pathogenicity, transmissibility, or immunogenicity of influenza in humans, we only report deep mutational scanning experiments that characterize the effects of HA mutations on binding to MHC-II from the tufted duck, a wild waterfowl species that is a natural host of H5 and H7 influenza^16,17^. We also survey the ability of HAs from human and avian influenza strains (taken from existing sequence databases) to bind to human MHC-II, but such characterization is confined to already known HA sequences, many from viruses which already infected humans. Therefore, our study provides insights into how influenza HA interacts with MHC-II from wild birds, and assesses the extent to which existing natural HAs can interact with human MHC-II, but does not report systematic characterization of HA mutations that might increase binding to human MHC-II.

There are potential benefits for vaccines and viral risk assessment of understanding how HA interacts with MHC-II, since (as mentioned in the Discussion of this paper) such binding could modulate the immunogenicity or host tropism of influenza virus. For instance, if HA’s binding to MHC-II dampens the immune response (a speculative but plausible idea), then HAs with reduced MHC-II binding could yield better vaccine immunogens. Because influenza viruses that bind MHC-II naturally infect humans (binding is present among some human endemic H1 strains and avian H5 strains that have caused recent human infections), safe studies to understand this binding have the potential to benefit human health.

### Data accessibility and computer code

Interactive visualizations and additional descriptions of the different deep mutational scanning datasets are available at the following sites:

- H5 HA deep mutational scanning: https://dms-vep.org/Flu-H5N1-American-Wigeon-2021-HA-tufted-duck-MHCII-DMS/
- H7 HA deep mutational scanning: https://dms-vep.org/Flu-H7-Anhui13-MHCII-binding/
- Tufted duck MHC-II deep mutational scanning: https://dms-vep.org/Tufted-duck-MHCII-DMS/

Computational pipelines used to analyse the deep mutational scanning data and the processed data are in the following GitHub repositories:

- H5 HA deep mutational scanning: https://github.com/dms-vep/Flu-H5N1-American-Wigeon-2021-HA-tufted-duck-MHCII-DMS/
- H7 HA deep mutational scanning: https://github.com/dms-vep/Flu-H7-Anhui13-MHCII-binding
- Tufted duck MHC-II deep mutational scanning:https://github.com/dms-vep/Tufted-duck-MHCII-DMS

A GitHub repository that builds the phylogenetic trees shown in this paper as well as additional phylogenetic trees that can be visualized interactively using Nextstrain is at https://github.com/jbloomlab/flu-ha-mhcii-usage-trees

The key numerical data on the effects of mutations after applying our recommended quality control filters are provided as both Supplementary Data and online spreadsheets:

- Effects of mutations in H5 HA on entry in different cell lines and binding to tufted duck MHC-II available as **Supplementary Data 1** and at https://github.com/dms-vep/Flu-H5N1-American-Wigeon-2021-HA-tufted-duck-MHCII-DMS/blob/master/results/summaries/tufted_duck_MHCII_binding.csv
- Effects of mutations in H7 HA on entry in different cell lines and binding to tufted duck MHC-II available as **Supplementary Data 2** and at https://github.com/dms-vep/Flu-H7-Anhui13-MHCII-binding/blob/master/results/summaries/tufted_duck_MHCII_binding.csv
- Effects of mutations in tufted duck MHC-II on binding to H5 HA available as **Supplementary Data 3** and at https://github.com/dms-vep/Tufted-duck-MHCII-DMS/blob/master/results/summaries/HA_binding.csv

A PDB file for the structural model of H5 HA in complex with tufted duck MHC-II is available at https://github.com/dms-vep/Flu-H5N1-American-Wigeon-2021-HA-tufted-duck-MHCII-DMS/blob/master/data/H5_MHCII_PDB/H5-4mut-MHCII-3mut-fit9-RBD.pdb and has been submitted to the Protein Data Bank with ID 36XU.

Plasmid maps of all constructs referred to in the methods are available at https://github.com/dms-vep/Flu-H5N1-American-Wigeon-2021-HA-tufted-duck-MHCII-DMS/tree/master/data/plasmid_maps

### Cell line maintenance

noSA, SA23, SA26 and 293T cell lines were maintained in D10 media (Dulbecco’s Modified Eagle Medium with 10% heat-inactivated fetal bovine serum, 2 mM l-glutamine, 100 U/mL penicillin, and 100 μg/mL streptomycin). Cell lines with rtTA expression were maintained in D10 media with tetracycline free FBS. Raji and SU-DHL-4 cells were grown in RPMI-1640 media (ATCC, 30-2001), supplemented with 10% heat-inactivated fetal bovine serum, 100 U/mL penicillin, and 100 μg/mL streptomycin. Calu-3 cells were grown in EMEM (ATCC, 30-2003) supplemented with 10% heat-inactivated fetal bovine serum, 100 U/mL penicillin, and 100 μg/mL streptomycin. To induce MHC-II expression, Calu-3 cells were treated with 500 U of IFN-γ (PeproTech, 300-02) for 48 hours.

### Cell lines expressing no sialic acid or only specific sialic acids

Generation of noSA, SA23 and SA26 cell lines is described in Narimatsu et al (2019)^11^ and Büll et al (2020)^12^. noSA cells are 293 cells that have had the genes for ST3GAL1/2/3/4/5/6 and ST6GAL1/2 sialyltransferases knocked-out. We have previously used these cells to study HA sialic acid preferences^10^. Note that while we refer noSA cells as not expressing sialic acids, they may retain some low levels of sialylated glycans produced by enzymes like ST6GALNAC/ST8SIA or other residual glycan pathways; either such residual low level sialic acid or some unknown alternative receptors likely explains why they can still be infected at very low but measurable efficiency by pseudoviruses expressing influenza HA. SA23 and SA26 are 293 cells that have ST3GAL3/4/6 and ST6GAL1/2 genes knocked out, and then further modified to overexpress either ST3GAL4 (SA23 cells) or ST6GAL1 (SA26 cells).

### Generation of MHC-II expressing cell lines for pseudovirus infections

We used noSA and 293T cells to generate cells overexpressing different MHC-II proteins. In all cases to generate MHC-II overexpressing cell lines, we used lentiviral transduction with the pHAGE2_EF1a_MHCII_IRES_Puro lentivirus backbone, which expresses MHC-II from an EF1a promoter and the puromycin resistance gene from an IRES. Note that all MHC-II expression constructs used in this manuscript have the same structure: MHC-II alpha chain, a T2A linker, and then MHC-II beta chain. See https://github.com/dms-vep/Flu-H5N1-American-Wigeon-2021-HA-tufted-duck-MHCII-DMS/tree/master/data/plasmid_maps for all plasmid maps.

To produce MHC-II overexpressing cell lines, we first generated VSV-G pseudotyped lentiviruses by transfecting 293T cells with lentiviral helper plasmids (BEI Resources: NR-52517, NR-52519, NR-52518), VSV-G expression plasmid, and pHAGE2_EF1a_MHCII_IRES_Puro backbone. 48 hours after transfection produced VSV-G pseudoviruses were clarified using a 0.45 μm SFCA syringe filter (Corning, Cat. No. 431220). 1 ml of the clarified supernatant was then used to transduce ∼1 million target cells (either 293T or noSA cells). As the lentivirus backbone contains no fluorescent marker we do not know the exact transcriptional units in 1 ml of used supernatant although we expect this to have been a high multiplicity of infection (MOI) transduction. Successfully transduced cells were then selected by performing 0.75 µg/ml puromycin selection until no cell death was observed (∼3 media changes). To measure human MHC-II expression in noSA, 293T and noSA-human-MHCII-1503 cells, we washed cells 3 times in phosphate-buffered saline with 1% bovine serum albumin (PBSB) buffer and incubated them on ice for 45 min with 1:100 dilution of BD OptiBuild™ RB744 Mouse Anti-Human HLA-DR antibody (BD Biosciences, 758140). We then performed 3 additional PBSB washes and antibody staining was detected by flow cytometry on a BD LSRFortessa X-50 machine.

### Production and titration of pseudoviruses expressing influenza HA

To produce lentiviral particles pseudotyped with HAs from H1, H2, H3, H5, H7 or H9 strains (pseudoviruses), we ordered codon optimized nucleotide sequences for each strain from Twist Bioscience. The protein sequences for all HAs are at https://github.com/dms-vep/Flu-H5N1-American-Wigeon-2021-HA-tufted-duck-MHCII-DMS/blob/master/data/HA_sequences.csv. For H5 HA pseudotyping, we selected a panel of strains that included representatives from each major clade of the GsGd-descended lineage, as well as several non-highly pathogenic H5 strains. For the smaller panels of H1, H2, and H3 HAs, we selected strains based on subtype-specific phylogenies, aiming to represent major lineages while including both human and non-human strains.

To produce the pseudoviruses, for each HA we transfected ∼1 million 293T cells with 1 µg of pHAGE-CMV-Luc2-IRES-ZsGreen-W-1270 lentiviral backbone, 0.25 µg each of lentiviral Gag/pol, Tat and Rev helper plasmids (BEI Resources: NR-52517, NR-52519, NR-52518), 0.1 µg of TMPRSS2 protease plasmid, 0.15 µg of neuraminidase (NA) expression plasmid, and 0.25 µg of HA expression plasmid. We used different NAs depending on a subtype: for H5 HA pseudoviruses we used N1 NA from the A/turkey/Indiana/22-003707-003/2022 strain, for H1 HA pseudoviruses we used N1 NA from the A/WSN/1933 strain, for H2 and H3 HA pseudoviruses we used N2 NA from the A/HongKong/1/1968, for H7 HA we used N9 NA from the A/Anhui/1/2013 strain, and for H9 HA we used N2 NA from the A/Hong Kong/33982/2009 strain. For transfection BioT transfection reagent (Bioland Scientific) was used according to manufacturer’s recommendations. 48 hours after transfection cell supernatant was clarified using a 0.45 μm SFCA syringe filter. Pseudoviruses were titrated on different cell lines that were plated with 2.5 µg/ml amphotericin B the day before, and titers were quantified as relative light units per µl (RLU/µl) using Bright-Glo Luciferase Assay System (Promega, E2610), as described previously^53^.

Note that H1, H2 and H3 HA pseudoviruses were produced and titrated in FBS-free neutralization assay media (NAM, Medium-199, 0.3% BSA, 100 U/mL penicillin, 100 μg/mL streptomycin, 100 μg/mL calcium chloride, and 25 mM HEPES), while the other subtypes were produced and titrated in D10 media.

To plot virus titers in relation to the phylogenetic trees as shown for instance in **Fig.1c**, we used the *tree-annotated-plot* package (v0.3.0, https://jbloomlab.github.io/tree-annotated-plot/).

### Generation of PB1 and MHC-II expressing cell lines for conditionally replicative influenza virus production and titration

For generation and/or infection of conditionally replicative H5N1 influenza viruses, we generated cells that express both PB1, which is required for virus replication, and tufted duck MHC-II. Generation of 293T-PB1 and MDCK-SIAT1-PB1-TMPRSS2 cells has been described previously^22,54^ and we overexpressed tufted duck MHC-II in these cells using the same protocol as described in ‘*generation of MHC-II expressing cell lines for pseudovirus infections’* section. To make noSA-PB1 and noSA-PB1-tufted-duck-MHCII cells we used noSA and noSA-tufted-duck-MHCII cells, respectively, and transduced them at high MOI with lentivirus made using pHAGE2_EF1A_PB1_CMV_HygroR lentiviral backbone, which contains PB1 gene from A/WSN/1933 strain under EF1α promoter and a hygromycin resistance gene under a CMV promoter. Note that this PB1 gene contains synonymous mutations that ablate expression of the PB1-F2 protein to avoid its toxic effect on the transduced cells, as described previously^22^. To select for successfully transduced cells we grew them in 100 ug/ml of hygromycin B until no cell death was observed (∼ 3 cell media changes).

### Conditionally replicative H5N1 influenza virus production and titration

Conditionally replicative influenza viruses were generated as shown in **Supplementary Figure 3a** and described previously^10,55,56^. In brief these are produced using pHW2000 reverse genetic plasmids^57^ that encode the coding sequence of H5 HA from A/American Wigeon/South Carolina/USDA-000345-001/2021 with polybasic cleavage site removed, N1 NA coding sequence is from A/turkey/Indiana/22-003707-003/2022, with the noncoding terminal regions for both HA and NA segments from the lab-adapted A/WSN/1933 strain. The coding sequence of PB1 gene is replaced by eGFP sequence and cloned into a unidirectional pHH-PB1flank-eGFP reverse genetics plasmid as described previously^22^. The other influenza segments (PB2, PA, NP, M, and NS) are expressed from the bidirectional reverse genetics plasmids for the lab-adapted A/WSN/1933 strain^57^. To generate conditionally replicative viruses, mixtures of either 5e5 of 293T-PB1 and 50,000 of MDCK-SIAT1-PB1-TMPRSS2 cells or 5e5 293T-PB1-tufted-duck-MHCII and 50,000 MDCK-SIAT1-PB1-TMPRSS2-tufted-duck-MHCII cell were plated. The next day cells were transfected with an equal-mass mixture of the reverse genetics plasmids encoding A/WSN/1933 PA, PB2, NP, M, NS^57^, pHH-PB1flank-eGFP, and the aforementioned reverse genetics plasmids encoding H5 HA and N1 NA plasmids. The next day, cell media was changed to NAM media and the virus-containing supernatant was collected 48 hours later. Images shown in **Supplementary Figure 6a** are from 48 hours post transfection. For titering of the resulting virus, supernatant was collected and cell debris was removed from the virus stock by a 10 min 1500 rcf spin at 4°C. To determine virus titers we plated noSA-PB1, noSA-PB1-tufted-duck-MHCII, 293T-PB1, and 293T-PB1-tufted-duck-MHCII cells and the next day we serially diluted virus stock on these cells. 16-20 hours post infection cells were washed and fixed with 4% paraformaldehyde solution and virus titers were determined by flow cytometry on BD LSRFortessa X-50 machine.

### Generation of tufted duck MHCII-VLPs

To generate virus-like particles covered in tufted duck MHC-II used for deep mutational scanning binding and neutralization measurements, 150 million 293T cells were plated into Falcon® 875cm^2^ 5-layer flasks (Corning, 353144). The next day these cells were transfected with 125 µg of lentiviral Gag/pol plasmid (BEI Resources: NR-52517) and 125 µg of HDM_tufted_duck_MHCII plasmid coding for tufted duck MHC-II. Note that Gag/pol plasmid codes for both lentiviral Gag and polymerase proteins in one overlapping reading frame, however, the polymerase protein itself does not serve any function in MHCII-VLP production. During transfection cell media was changed to influenza growth media (Opti-MEM supplemented with 0.3% bovine serum albumin, 100 U/mL penicillin, and 100 µg/mL streptomycin) to minimize amount of sialic acids in cell supernatant. 48-72 hours post transfection the MHCII-VLP-containing cell supernatant was clarified using SFCA, 0.45 µm filter (Thermo Scientific, 1564045) and concentrated by ultracentrifugation over 20% sucrose cushion (made in 1x tris-buffered saline) by spinning at 100,000g for 1 hour using Beckman Coulter SW 32 Ti rotor. MHCII-VLP pellets were resuspended in 1x tris-buffered saline and frozen at -80°C until further use.

### Inverted pseudotyping of lentiviral particles with MHC-II

Expression plasmids with different MHC-II genes used for inverted pseudotyping in this manuscript are available at https://github.com/dms-vep/Flu-H5N1-American-Wigeon-2021-HA-tufted-duck-MHCII-DMS/tree/master/data/plasmid_maps. To pseudotype lentiviral particles with MHC-II, we transfected ∼1 million 293T cells with 1 µg of pHAGE-CMV-Luc2-IRES-ZsGreen-W-1270 lentiviral backbone, 0.25 µg each of lentiviral Gag/pol, Tat and Rev helper plasmids (BEI Resources: NR-52517, NR-52519, NR-52518), 0.15 µg of N1 NA expression plasmid, and 0.25 µg of MHC-II expression plasmid. 48 hours post transfection cell supernatant was clarified using 0.45 μm SFCA syringe filter (Corning, Cat. No. 431220). Viral titers were determined by titration on 293T-H5-HA cell lines that were plated a day before in the presence of 1 µg/ml of doxycycline and virus titers were read out as relative light units per µl (RLU/µl) using Bright-Glo Luciferase Assay System (Promega, E2610), as described previously^53^.

Note that we co-transfect NA expression plasmids during MHC-II pseudovirus production to cleave sialic acids from lentiviral particles. This is important because we found that lentiviral particles made without any MHC-II (bald particles) could efficiently enter HA expressing target cells when no NA was included, likely due to the binding of sialic acids present on lentiviral particles to the HA expressed on target cells. This effect was most notable on target cells that express high levels of HA (like our 293T-H5-HA cells), likely because of high avidity required for sialic acid mediated entry.

### H5 HA expressing cell lines for infection with MHC-II pseudoviruses

293T-H5-HA cells, which we used as target cells for MHC-II pseudovirus infections, are 293T-rtTA^9^ cells (see *production of tufted duck MHC-II deep mutational scanning plasmid libraries* section) that were first transduced with a doxycycline-inducible lentivirus backbone expressing a lentiviral tat gene (pHAGE2_TRE3G_tat), clonally expanded, and then transduced at high MOI with a lentivirus backbone expressing H5 HA from A/American Wigeon/South Carolina/USDA-000345-001/2021 strain. The specific lentivirus backbone used for HA expression, pH2rU3_ForInd_H5_H5N1_turkey_Indiana_genscript_T7_CMV_RFP657T2APurR, has repaired U3 region in the lentiviral LTR enabling Tat-mediated transcription^9^, a TRE3G promoter enabling inducible HA expression, and an RFP657 fluorescent protein linked by T2A to puromycin resistance gene under a CMV promoter. Successfully trans-duced cells were selected by puromycin. The reason for this design is that we wanted to have a cell line with inducible HA expression to limit cell toxicity and potential loss of HA expression over time, however, the TRE3G promoter that allows doxycycline-inducible HA expression is not as strong a promoter as the lentiviral tat promoter. Therefore, we used cells that have inducible tat expression to further increase expression of HA from the lentiviral backbone. In these cells addition of doxycycline to 293T-H5-HA cells induces expression of both tat and HA and the expressed tat then further increases expression of the HA. Note that expression of HA in these cells is so high that even bald lentiviral particles can enter these cells with high titers using sialic acids on their surface, so expression of NA on pseudoviruses is required to avoid sialic acid mediated entry of pseudoviral particles (see *Inverted pseudotyping of lentiviral particles with MHC-II* for details).

### H5 and H7 HA deep mutational scanning libraries

The H5 HA deep mutational scanning libraries used in this study were described previously in Dadonaite et al (2024)^10^. These libraries cover almost every possible amino-acid mutation in H5 HA from the A/American Wigeon/South Carolina/USDA-000345-001/2021 strain. The H7 HA deep mutational scanning libraries used in this study were described previously in Ahn et al (2026)^33^, and cover almost every possible amino-acid mutation in the ectodomain of the HA from the A/Anhui/1/2013 strain. The pseudoviruses in both libraries consist of lentiviral particles expressing both HA and NA proteins on their surface, with the HAs being different representatives of the mutant library with a genotype-phenotype link between the virion surface HA protein and a barcode-linked HA gene in the lentiviral genome, and the NA being the unmodified protein from the homologous strain but no NA gene in the lentiviral genome. Note that there are two independent mutant libraries for both the H5 and H7 HA pseudoviruses, and all experiments in this study performed independent experiments with each of the two libraries and report the mean measurement across the two libraries.

### Measurement of differences in HA mutation effects on cell entry in MHC-II versus sialic acid expressing cells by deep mutational scanning

To calculate the differences in mutation entry effects between sialic acid and tufted duck MHC-II expressing cells for H5 and H7 HAs, we first measured how mutations affect entry in these cell lines. For H5 deep mutational scanning libraries, we measured entry effects in SA23, SA26, an equal mixture of SA23 and SA26 cells, and noSA-tufted-duck-MHC-II cell lines. For H7 deep mutational scanning libraries, we measured entry effects in 293T and noSA-tufted-duck-MHC-II cell lines. These experiments were performed as described before^9,10,33^ with a few modifications. For each cell line we plated 3 million cells in duplicate a day before infection. One replicate was plated with 2.5 µg/ml amphotericin B as amphotericin B increases HA pseudovirus infection^10^ but the other replicate had no amphotericin B added because it decreases VSV-G pseudovirus infection. The day of infection, we changed cell media to influenza growth media. For both H5 and H7 HA deep mutational scanning libraries, we used 6 million transcription units of HA pseudovirus and infected cells that were plated with amphotericin B. Note that library pseudoviruses used to infect cell lines that express sialic acids were preincubated for 20 min on ice with 1uM oseltamivir in order to inhibit NA to prevent it from cleaving sialic acids on the target cells. No oseltamivir was used on MHC-II expressing cell lines, thereby enabling NA to cleave any residual sialic acids on noSA-tufted-duck-MHC-II cells and so reduce background non-MHC-II–mediated entry. For the control VSV-G pseudotyped H5 and H7 pseudoviruses libraries used to normalize for frequencies of different variants^9^, we used 10 million transcription units and infected cells plated without amphotericin B. 15-20 hours post infection non-integrated lentiviral genomes were recovered using QIAprep Spin Miniprep Kit and amplicons were prepared for Illumina sequencing as described previously^9^. We used dual indexing for each sample to avoid index hopping. Sequencing was performed on NextSeq 2000 and NovaSeq X Plus platforms. All cell entry effect measurements were performed in biological duplicate using the two independent deep mutational scanning libraries for both H5 and H7 HAs.

To quantify the effects of each mutation on cell entry, we first calculated a log enrichment ratio (functional score) for each barcoded HA variant as *log*_2_ ([*n*^*v*^_*post*_ / *n*^*wt*^_*post*_] /[*n*^*v*^_*pre*_/ *n*^*wt*^_*pre*_], where *n*^*v*^_*post*_ is the count of variant v in the HA-pseudotyped infection , *n*^*v*^_*pre*_ is the count of variant v in the VSV-G-pseudotyped infection, and *n*^*wt/*^_*post*_ and *n*^*wt/*^_*pre*_are the counts for unmutated variants in the HA pseudotyped versus VSV-G pseudotyped infections. We then jointly analyzed the functional scores for singly and multiple mutated variants using a sigmoid global epistasis function^58^ to estimate single mutation effects. Examples of the global epistasis fitting can be seen:

- For H5 HA libraries: https://dms-vep.org/Flu-H5N1-American-Wigeon-2021-HA-tufted-duck-MHCII-DMS/notebooks/func_effects_global_epistasis_Lib1-251017-293-noSA-duck-MHCII.html
- For H7 HA libraries: https://dms-vep.org/Flu-H7-Anhui13-MHCII-binding/note-books/func_effects_global_epistasis_LibA-251202-H7_293T.html

See appendices in the homepages linked in the *Data accessibility and computer code* section for comparable notebooks showing fitting for all replicates on all cells tested. We then averaged the mutation effects for the replicate libraries and report the average effect across libraries. We also applied quality-control filters to remove measurements for mutations not observed in at least two barcoded variants or mutations with high standard deviation in measured effects across the replicate libraries.

To calculate the differences in effects of mutations in MHC-II versus sialic acid expressing cells, we subtracted cell entry effects measured on noSA-tufted-duck-MHC-II cells from effects measured on sialic acid expressing cells. An example notebook performing this calculation is shown at https://dms-vep.org/Flu-H5N1-American-Wigeon-2021-HA-tufted-duck-MHCII-DMS/notebooks/func_effect_diffs_tufted-duck-MHCII_vs_SA26-SA23_entry.html and other comparable notebooks are shown in the appendices of the homepages linked in the data availability statement.

### Measurement of effects of HA mutations on tufted duck MHCII-VLP binding by deep mutational scanning

To measure how H5 and H7 HA mutations affect binding to MHCII-VLPs, we incubated 1.5 million transcription units of H5 and H7 HA deep mutational scanning libraries with multiple dilutions of concentrated MHCII-VLPs (see *generation of tufted duck MHCII-VLPs*). We aimed to cover most of the neutralization range with the chosen dilutions in order to identify mutations that both increased and decreased MHCII-VLP binding. The dilutions selected neutralized deep mutational scanning libraries with inhibitory concentrations in the range of IC6 to IC95. However, because the MHCII-VLP preparations likely contained heterogeneous particles as well as some free MHC-II, achieving consistent neutralization across libraries was not always possible. The libraries were incubated with MHCII-VLP dilutions for 30 min at 37°C and the virus-VLP mixtures were used to infect 1 million 293T cells that were plated 24 hours earlier. 15-20 hours post infection non-integrated lentiviral genomes were recovered using QIAprep Spin Miniprep Kit. Note that a non-neutralizable DNA standard was added to the cell lysis step (P2 buffer) to the amount equal to ∼2% of the total recovered non-integrated lentiviral genomes in order to act as a non-neutralizable standard to convert barcode counts into absolute pseudovirus infectivity retained using a previously described approach^9^. Non-integrated genome recovery and subsequent amplicon library preparation for Illumina sequencing was described previously^10,33^. These experiments were performed in biological duplicates using the two independent deep mutational scanning libraries.

Analysis of mutation effects on MHCII-VLP binding was done by converting variant barcode counts into fraction infectivity retained using the non-neutralizable DNA standard, and then analyzing these fraction infectivities using *polyclonal*^59^ package version 6.17 (https://jbloomlab.github.io/polyclonal/) integrated in dms-vep-pipeline-3 version 3.34 (https://github.com/dms-vep/dms-vep-pipeline-3) to estimate the effects of individual mutations. Example of the model fitting can be seen:

- For H5 HA deep mutational scanning libraries https://dms-vep.org/Flu-H5N1-American-Wigeon-2021-HA-tufted-duck-MHCII-DMS/notebooks/fit_escape_MHCII_binding_Lib2-251218-duck_MHCII.html
- For H7 HA deep mutational scanning libraries https://dms-vep.org/Flu-H7-Anhui13-MHCII-binding/notebooks/fit_escape_MHCII_binding_B-251218-duck_MHCII.html

See appendices in the homepages linked in the *Data accessibility and computer code* section for comparable notebooks showing fitting for all library replicates.

### Validation of HA mutation effects on binding to tufted duck MHCII-VLP as assessed by HA pseudovirus neutralization

Expression plasmids with codon optimized unmutated and variant H5 HA sequences used for MHCII-VLP binding validation are provided at https://github.com/dms-vep/Flu-H-5-N1-American-Wigeon-2021-HA-tufted-duck-MHCII-DMS/-t-ree/master/data/plasmid_maps. We generated lentiviral particles pseudotyped with unmutated or variant HAs by transfecting ∼1 million 293T cells with 1 µg of pHAGE-CMV-Luc2-IRES-ZsGreen-W-1270 lentiviral backbone, 0.25 µg each of lentiviral Gag/pol, Tat and Rev helper plasmids, 0.15 µg of neuraminidase expression plasmid, and 0.25 µg of HA expression plasmid. 48 hours post transfection cell supernatant was clarified using 0.45 μm SFCA syringe filter. Viruses were titrated on 293T cells that were plated with 2.5 µg/ml amphotericin B the day before and virus titers were read out as relative light units per µl (RLU/µl) using Bright-Glo Luciferase Assay System. To perform neutralization assays, unmutated or variant pseudoviruses were incubated with a series of 8 different 3-fold serial dilutions of concentrated MHCII-VLPs for 1 hour at 37°C and subsequently used to infect 293T cells that were plated with 2.5 µg/ml amphotericin B a day before. 48 hours after infection virus titers at each dilution were read out using Bright-Glo Luciferase Assay System and fraction infectivity was calculated using controls that were not incubated with MHCII-VLP. The neutcurve software^60^ (https://jbloomlab.github.io/neutcurve/) was used to fit Hill curves to the fraction infectivity data.

### Production of tufted duck MHC-II deep mutational scanning plasmid libraries by Golden Gate assembly

Tufted duck MHC-II libraries were designed to include every possible amino acid mutation in the ectodomains of tufted duck MHC-II alpha and beta chains. Signal peptides, transmembrane domains, cytoplasmic tails, and the T2A linker connecting the two chains were not mutagenised. We intentionally included stop codons in the first 20 amino acid positions in the alpha chain ectodomain as controls (these mutations are expected to be highly deleterious). The sequence of the unmutated codon optimized MHC-II cloned into lentiviral vector used for deep mutational scanning is available at https://github.com/dms-vep/Tufted-duck-MHCII-DMS/blob/master/library_design/5657_pHrU3_ForInd-Extgag_duck_MHCII_opt4.gb. Note that the tufted duck MHC-II codon optimization was specifically chosen for its ability to produce high-titer pseudoviruses from single-copy integrated lentivirus backbones^9^.

The plasmid library for the tufted duck deep mutational scanning was created using Golden Gate assembly^61–63^ (GGA) using a previously described strategy^64,65^, with additional modifications. First, we divided the tufted duck MHC-II sequence into 6 tiles for GGA, which together covered the full mutagenized MHC-II region. The sequence for each tile fragment is available at https://github.com/dms-vep/Tufted-duck-MHCII-DMS/blob/master/library_design/tufted_duck_MHCII_assembly_fragments.csv. In addition, to the ends of each tile we added flanking BsmBI sites and unique primer sequences that were later used to amplify each tile fragment from a single oligo pool. We then computationally designed every mutation we wanted to introduce into the library as an individual DNA fragment in an oligo pool. For the code used to do this, see https://github.com/dms-vep/Tufted-duck-MHCII-DMS/blob/master/library_design/gga_codon_muts_oligo_design.py. This script takes a list of intended mutations and tile sequences and creates a fasta file that contains oligo sequences with all desired mutations. The output fasta file the script generates can be used to directly order oligo pools from Twist Biosciences and it is available here https://github.com/dms-vep/Tufted-duck-MHCII-DMS/blob/master/library_design/GGA_oPool.fa

We used the synthesized oligo pool as a template to separately PCR amplify each of the six mutated tile pools. We did so by using primers annealing to the unique sequences designed to flank each tile; these primers are available at https://github.com/dms-vep/Tufted-duck-MHCII-DMS/blob/master/library_design/primers.csv. The PCR reactions to amplify each tile used 1ng of oligo pool template, 0.3 µM of forward and reverse primer, and 1x KOD Hot Start Master Mix (Sigma-Aldrich, Cat. No. 71842). Each PCR reaction was started at 95°C for 2 min and then went through 23 cycles of 95°C for 20 s, 62°C for 10 s, 68°C for 25 s. The PCR products were purified with Ampure XP beads using 1:3 DNA to bead ratio. For each tile, we created a destination vector into which the amplified tile pools were assembled. The destination vectors consisted of pGGAselect vector (NEB, E1602S; cloned to have ampicillin resistance) into which we cloned MHC-II sequences where each tile region is flanked by BsmBI sites at both ends. In addition, the tile region between BsmBI sites in the destination vector was truncated by around ∼200nt so that in case the GGA reaction does not fully remove the uncut destination vector it would be removed during a gel size selection step. Plasmids maps for destination vectors are available at https://github.com/dms-vep/Tufted-duck-MHCII-DMS/tree/master/library_design/destination_vectors. We performed GGA assembly for each amplified tile pool and destination vector using NEBridge Golden Gate Assembly Kit (BsmBI-V2) according to manufacturer’s instructions. 100 fmol of tile pool and 50 fmol of destination vector was used and GGA reaction was incubated at 42°C for 1 hour followed by 60°C, for 5 minutes. Next each GGA reaction was purified using Ampure XP beads, eluted in 20 µl of water and 1 µl of eluted reaction was electroporated into NEB® 10-beta Electrocompetent E. coli cells (C3020K) according to the manufacturer’s instructions. Electroporated cells were outgrown overnight in ampicillin-containing LB media and next day assembled plasmids were recovered using QIAprep Spin Miniprep Kit. Note that >1 million colony forming units were recovered for each tile. Full length mutagenised MHC-II sequence was then amplified and barcoded from each cloned destination vector. We performed PCR on each cloned tile vector using KOD Hot Start Master Mix, 10 ng of plasmid library and 0.3 µM of forward (5′-gcacgcgCAGCCGAGCCACATCGCTCA-3′) and reverse (5′-gcggaactcc actaggaacatttctctctcgaaTCTAGANNNNNNNNNNNNNNNNAGATCGGAAGAGCGT CGTGTAGGGAAAGAG-3′) primers, the latter primer contained a 16 nt barcode. Correct PCR size products were then size selected on an agarose gel and purified using Ampure XP beads. At this point all six barcoded products were pooled equimorally, accounting for the number of mutations introduced per tile. We created two independently pooled reactions, which were then used to make two biological library replicates (Lib-1 and Lib-2). The pooled MHC-II libraries were then HiFi cloned into pre-digested lentiviral backbone used for deep mutational scanning^64,66^, electroporated into NEB® 10-beta Electrocompetent E. coli cells and outgrown overnight in ampicillin-containing LB media. We performed 5-7 electroporation reactions per library to recover >7 million colonies per library, which is required in order to maintain high barcode diversity and to avoid barcode duplication in later library production steps.

### Production of cell stored tufted duck MHC-II deep mutational scanning libraries

To produce cell-stored deep mutational scanning libraries for the pseudoviruses encoding tufted duck MHC-II, we first made a 293T-rtTA-tdCD74 cell line in which the libraries were stored. To do so we used 293T-rtTA cells (a 293T clone we have previously found to yield particularly high pseudovirus titers^9^ that has been engineered to overexpress reverse tetracycline-controlled transactivator, rtTA). For tufted duck MHC-II libraries, we further modified 293T-rtTA cells to also overexpress an invariant chain from a tufted duck (tdCD74, Genbank accession XP_032052821.1). The reason that we expressed tdCD74 is that MHC-II overexpression in cells can negatively affect pseudovirus titers, and coexpression of CD74 can improve lentivirus titers by around 3-5 fold (data not shown), perhaps due to reduced proteostasis cell stress due to improved MHC-II trafficking. To produce 293T-rtTA-tdCD74 cells, we used the same protocol as for MHC-II overexpressing cell line production except pHAGE2_EF1A_tufted_duck_CD74_CMV_HygroR lentiviral backbone was used. To select for successfully transduced 293T-rtTA cells we passaged them in 100 ug/ml of hygromycin B until no cell death was observed (∼3 cell media changes).

To produce cell-stored tufted duck MHC-II deep mutational scanning libraries, we followed our previously described method^9^. First, we produced VSV-G pseudotyped lentiviral particles carrying lentiviral backbones with mutagenised tufted duck MHC-II followed by a 16 nt barcode. To do so, we transfected 20 million 293T cells with 20 µg of MHC-II library, 5 µg of each lentiviral helper plasmid, 5 µg of tdCD74, and 5 µg VSV-G expression plasmid. We co-transfected tdCD74 at this stage to avoid enriching the libraries with non-functional MHC-II variants since as mentioned above, MHC-II overexpression affects general protein expression but tdCD74 can ameliorate this effect. At 48 hours after transfection, we collected and purified the virus-containing supernatant using a 0.45 μm SFCA syringe filter. We then transduced 293T-rtTA-tdCD74 cells with the VSV-G viruses at low MOI (<0.01), so that on average no more than one lentiviral backbone was integrated into each cell. Successfully transduced cells were selected using puromycin and stored at -80 °C long-term. Each of these cells then stores a uniquely barcoded tufted duck MHC-II genome in the form of an integrated proviral vector.

### Long-read PacBio sequencing for variant-barcode linkage

Long read sequencing was performed on lentiviral genomes to link mutations present in MHC-II with the associated barcodes as described previously^9^. Namely, VSV-G pseudo-typed lentiviral particles were produced from cell-stored libraries to capture the full set of variants present in MHC-II libraries independent of MHC-II’s ability to fold and mediate entry. 10 million transcriptional units of these VSV-G pseudotyped particles were then used to infect 3 million 293T cells and the next day non-integrated viral genomes were recovered using QIAprep Spin Miniprep Kit. The full length MHC-II and associated barcode sequences were amplified in two rounds of PCR, keeping the number of PCR cycles low to minimize PCR strand exchange. Amplified genomes were then prepared for PacBio sequencing using SMRTbell prep kit 3.0 (PacBio, 102-182-700), according to the manufacturer’s instructions. The prepared long-read libraries were sequenced on PacBio Revio system.

The resulting PacBio circular consensus sequences (CCSs) were analyzed using *alignparse*^*67*^ (https://github.com/jbloomlab/alignparse) as integrated into version 3.34.0 of *dms-vep-pipeline-3* (https://github.com/dms-vep/dms-vep-pipeline-3). Notebooks detailing the analysis can be found in the “Barcode to codon-variant lookup table” section of the appendix of the homepage for the tufted duck MHC-II deep mutational scanning page (https://dms-vep.org/Tufted-duck-MHCII-DMS/appendix.html).

The total number of variants in the tufted MHC-II libraries was 96,734 and 104,597 for Lib-1 and Lib-2, respectively. The average number of mutations per variant was 1.32 and 1.29 for Lib-1 and Lib-2, respectively. The variant-barcode look-up table for both biological library replicates made using at least 2 CCSs is available here https://github.com/dms-vep/Tufted-duck-MHCII-DMS/blob/master/results/variants/codon_variants.csv

### Measuring effects of mutations in tufted duck MHC-II on inverted pseudovirus entry in 293T-H5-HA cells by deep mutational scanning

To measure how mutations in tufted duck MHC-II affect cell entry, we first produced MHC-II or VSV-G pseudotyped lentiviral particles from the cell-stored libraries. To produce VSV-G pseudotyped library pseudoviruses, we plated library cells in Falcon® 875cm^2^ 5-layer flasks and the next day transfected them with 50 µg of each lentiviral helper plasmid, 23 µg of VSV-G plasmid, and 18 µg of tdCD74 plasmid. 48 hours after transfection the cell supernatant was clarified using SFCA, 0.45 µm filter and concentrated using Pierce Protein Concentrator (ThermoFisher, 88537). MHC-II pseudotyped viruses were produced the same way except library cells were plated with 1 µg/ml of doxycycline to induce MHC-II expression and VSV-G plasmid was excluded from the transfection.

To measure how mutations in tufted duck MHC-II affect entry in 293T-H5-HA cells, we plated 3 million 293T-H5-HA and 293T cells a day before infection. 293T-H5-HA cells were plated in the presence of 1 µg/ml of doxycycline to induce HA expression. The 293T-H5-HA cells were then infected with 5 million transcription units of MHC-II pseudotyped library viruses and 293T cells were infected with 30 million transcription units of VSV-G pseudotyped viruses. 15-20 hours post infection non-integrated lentiviral genomes were recovered using QIAprep Spin Miniprep Kit and amplicons were prepared for Illumina sequencing as described previously^9^.

Effects of individual mutations on entry into 293T-H5-HA cells were calculated as described in *Measurement of differences in HA mutation effects on cell entry in MHC-II versus sialic acid expressing cells by deep mutational scanning* section and previously^9^. We use these measured cell entry scores as a filter for MHC-II HA binding data (see below) as our binding measurements rely on the ability of MHC-II pseudoviruses to at least enter the cells, and low entry scores indicate non-functional MHC-II for which accurate binding measurement cannot be made.

### Measuring effects of mutations in tufted duck MHC-II on H5 HA binding by deep mutational scanning

To measure how mutations in tufted duck MHC-II affect binding to H5 HA from A/American Wigeon/South Carolina/USDA-000345-001/2021 strain, we performed neutralization experiments of the MHC-II pseudovirus libraries with soluble HA trimer. MHC-II mutants that bind more strongly to HA are neutralized more potently by soluble HA trimer. For these experiments, 1.5 million transcriptional units of MHC-II pseudovirus libraries were incubated with increasing concentrations of soluble HA trimer for 45 min at 37°C. The concentrations of the trimer used were 4.83, 9.67, 19.34, 48.35, and 120.87 µg/ml, which reached between IC30 and IC89 neutralization of the library pseudovirus. 1 million 293T-H5-HA cells pre-plated with doxycycline were then infected with pseudovirus-HA mixtures. 15-20 hours post infection, non-integrated lentiviral genomes were recovered using QIAprep Spin Miniprep Kits and amplicons were prepared for Illumina sequencing as described previously^9^. Note that a non-neutralizable DNA standard was added to the cell lysis step (P2 buffer) to the amount equal to ∼1% of the total recovered non-integrated lentiviral genomes. Fraction neutralization values for each variant were determined relative to the non-neutralizable DNA standard and mutation effects on HA binding were calculated using a biophysical model implemented in *polyclonal*^*59*^ package version 6.17 (https://jbloomlab.github.io/polyclonal/) integrated in dms-vep-pipeline-3 version 3.34 (https://github.com/dms-vep/dms-vep-pipeline-3). An example of the model fitting can be seen https://dms-vep.org/Tufted-duck-MHCII-DMS/notebooks/fit_escape_HA_binding_Lib1-260529-HA_binding.html.

### Protein Expression and Protein Purification

All recombinant proteins used in this study were codon-optimized for human cell expression and made in the CMV/R vector^68^ by Genscript with a C-terminal hexahistidine affinity tag. PEI MAX was used for transient transfection of HEK293F cells. All H5 HA constructs were co-transfected with N1 NA at 20:1 HA:NA to facilitate secretion of soluble HA from producing cells. After four days, mammalian cell supernatants were clarified via centrifugation and filtration. All proteins were purified using ion-metal affinity chromatography. 1 mL of Ni2+-sepharose Excel was added per 100 mL clarified supernatant along with 5 mL of 1 M Tris, pH 8.0 and 7 mL of 5 M NaCl and left to batch bind while shaking at room temperature for at least 30 min. Resin was then collected in a gravity column, washed with 5 column volumes of 50 mM Tris, pH 8.0, 500 mM NaCl, 20 mM imidazole, and protein was eluted using 50 mM Tris, pH 8.0, 500 mM NaCl, 300 mM imidazole. Further purification was done using SEC on a Superdex 200 Increase 10/300 gel filtration column equilibrated in 25 mM Tris, pH 8.0, 150 mM NaCl, 5% glycerol. H5 HA was biotinylated via a C-terminal AviTag using a BirA biotin-protein ligase standard reaction kit (Avidity). Biotinylated H5 HA was then purified again by SEC as described above.

### Protein expression constructs

HA protein construct used in this study came from the A/American Wigeon/South Carolina/USDA-000345-001/2021 strain. NA protein constructs used in this study came from the A/turkey/Indiana/22-003707-003/2022 H5N1 strain. All H5 HA constructs were composed of ectodomains with three known stabilizing mutations (H355W, K380M, E432L)^31,69^ (**Supplementary Data 4**) and hexahistidine tags for affinity purification. H5 HA trimers for BLI were fused to the foldon trimerization domain^70^ and also contained an AviTag for *in vitro* biotinylation. H5 HA trimers for EM were fused to the nanoparticle trimer component I53-50A^71^. Tufted duck MHC-II dimer was made using a single plasmid with a T2A peptide sequence between the alpha and beta chains, the transmembrane domains replaced with a soluble dimerization domain^72^ and a hexahistidine tag.

### Bio-layer interferometry (BLI)

BLI was carried out using an Octet Red 96 system, at 25°C with 1000 rpm shaking. Biotinylated H5 HA was diluted in kinetics buffer (PBS with 0.5% bovine serum albumin and 0.01% Tween) to a final concentration of 20 μg/mL before loading onto Streptavidin biosensors (Sartorius) for 300 seconds. MHC-II concentrations were 2.3 μM, 1.15 μM, 0.58 μM, 0.29 μM, and 0.14 μM for both assays with H5 HAs against wild type MHC-II and 4 μM, 2 μM, 1 μM, 0.5 μM, 0.25 μM, and 0.125 μM for testing the H5 HA variant against the MHC-II variant in kinetics buffer, and their association was measured for 300 seconds, followed by dissociation for 300 seconds in kinetics buffer alone.

### Negative Stain Electron Microscopy (nsEM)

30 μM MHC-II was mixed with 6 μM H5 HA for at least 1 hour at room temperature. MHCII-H5 HA mixtures were diluted to 10 µg/mL and immediately applied to glow-discharged 400-mesh carbon-coated grids (Electron Microscopy Sciences) and stained with 2% (wt/vol) uranyl formate. Data were collected using EPU 2.0 on a 120 kV Talos L120C transmission electron microscope (Thermo Scientific) with a BM-Ceta camera. CryoSPARC^73^ was used for CTF correction, particle picking using the blob picker, particle extraction, and 2D classification. We calculated the approximate percentage of H5 HA bound to MHC-II by selecting all H5 HA 2D classes (both bound and unbound) and then summing the percentages of particles that made up classes bound to MHC-II relative to this total population

### Cryo-EM sample preparation, data collection and processing

Cryo-EM protein complexes of H5 HA containing I80L+A137S+T219E+R227G with tufted duck MHC-II containing G23M, K42I and Q65F mutations were prepared as described above for nsEM, with the addition of 0.1% w/v OBG detergent on glow-discharged Au QUANTIFOIL 1.2/1.3 holey-carbon grids (Electron Microscopy Sciences). Protein sample at 2 mg/ml was applied to the grid, incubated for 30 seconds, and then blotted using Whatman filter paper; this process was performed twice, followed by a third application of sample and final blotting and plunge freezing in liquid-ethane using a Vitrobot (Ther-moFisher). Data were acquired on a Glacios microscope (ThermoFisher) operating at 200 kV equipped with a K3 Summit Direct Detector (Gatan Inc.). Movies were acquired in counting mode, collecting 100 frames with a total dose of 54 electrons/Å2, with a defocus range of −0.5 to −2 µm (**Supplementary Data 5**). Automated data collection was carried out using SerialEM^74^ at a nominal magnification of 45,000× with a pixel size of 0.866 Å. All data processing was carried out using cryoSPARC^73^. Movie frame alignment and dose-weighting summation were performed using Patch Motion, and CTF estimation was done using patch CTF. Particles were first picked using Blob picker and these were used to generate initial 2D averages. Three more rounds of 2D classification were performed to select all well-aligned, single HA-MHCII complex views. An initial model was generated by ab-initio reconstruction with C1 symmetry, followed by an initial homogeneous 3D refinement with C3 symmetry. Symmetry expansion applying C3 symmetry was then performed, followed by 3D classification of single HA-MHCII protomers with a solvent mask around the entire trimeric HA-MHCII complex and a focus mask around a single HA-MHCII monomer. Only 3D classes with strong MHC-II density observed were selected for a final local refinement with C1 symmetry using the aforementioned focus mask. This resulted in a 3.4 Å reconstruction that was used to relax an H5 HA-MHCII model.

### Model building and refinement

Residues 11-329 in a single HA1 chain from a previously solved H5 HA cryo-EM structure (9DWE) and an AlphaFold3^75^ model of the tufted duck MHC-II dimer were docked into the cryo-EM map as rigid bodies using ChimeraX^76^. ChimeraX was used to make a sequence-accurate initial model of this H5 HA variant. A single N-acetylglucosamine at each non-clashing glycan sequon was added using GlycoShape^77^. A refinement of one H5 HA-MHCII protomer into the cryo-EM density was performed with ISOLDE^78^. A final real-space refinement with grid searching, Ramachandran and rotamer restraints all turned off, and ‘use starting model as reference’ turned on, was then performed in Phenix^79^. Flexible loops at the outer edge of the MHC-II molecule were not well resolved in the cryo-EM density so these loops were not modeled.

## Supplementary Data

### Supplementary Data 1. Effects of mutations in H5 HA on entry in different cell lines and binding to tufted duck MHCII

A version of this file is also available at https://github.com/dms-vep/Flu-H5N1-American-Wigeon-2021-HA-tufted-duck-MHCII-DMS/blob/master/results/summaries/tufted_duck_MHCII_binding.csv

### Supplementary Data 2. Effects of mutations in H7 HA on entry in different cell lines and binding to tufted duck MHCII

A version of this file is also available at https://github.com/dms-vep/Flu-H7-Anhui13-MHCII-binding/blob/master/results/summaries/tufted_duck_MHCII_binding.csv

### Supplementary Data 3. Effects of mutations in tufted duck MHC-II on binding to H5 HA

A version of this file is also available at https://github.com/dms-vep/Tufted-duck-MHCII-DMS/blob/master/results/summaries/HA_binding.csv

### Supplementary Data 4. HA, tufted duck MHC-II and NA protein sequences used for recombinant protein expression for structural analyses

A version of this file is also available at https://github.com/dms-vep/Flu-H5N1-American-Wigeon-2021-HA-tufted-duck-MHCII-DMS/blob/master/data/H5_MHCII_PDB/constructs.csv

### Supplementary Data 5. Cryo-EM data collection and refinement statistics

