## Supplementary Data 5 for "Defining the molecular interaction between influenza hemagglutinin and MHC-II"

### **Supplementary Data 5. Cryo-EM data collection and refinement statistics.**

| H5 HA trimer variant binding to tufted duck MHC-II variant  PDB 9PR3  EMD 71796 | |
| --- | --- |
| **Data collection and processing** | |
| Magnification (×) | 45,000 |
| Voltage (kV) | 200 |
| Electron exposure (e^–^/Å^2^) | 55 |
| Defocus range (μm) | -0.5 - -1.5 |
| Pixel size (Å) | 0.866 |
| Symmetry imposed | C1 |
| Final particle images (no.) | 104,785 |
| Map resolution (Å) | 3.4 |
| FSC threshold | 0.143 |
| Map sharpening *B* factor (Å^2^) | -90 |
| **Validation** | |
| MolProbity score | 0.77 |
| Clashscore | 0 |
| Poor rotamers (%) | 0.2 |
| Ramachandran plot |  |
| Favored (%) | 96.0 |
| Allowed (%) | 3.8 |
| Disallowed (%) | 0.2 |
